# ROCK and the actomyosin network control biomineral growth and morphology during sea urchin skeletogenesis

**DOI:** 10.1101/2022.12.21.521381

**Authors:** Eman Hijaze, Tsvia Gildor, Ronald Seidel, Majed Layous, Mark Winter, Luca Bertinetti, Yael Politi, Smadar Ben-Tabou de-Leon

**Author notes:** Correspondence: Smadar Ben-Tabou de-Leon.

## Abstract

Biomineralization had apparently evolved independently in different phyla, using distinct minerals, organic scaffolds and gene regulatory networks (GRNs). However, diverse eukaryotes from unicellular organisms, through echinoderms to vertebrates, use the actomyosin network during biomineralization. Specifically, the actomyosin remodeling protein, Rho-associated coiled-coil kinase (ROCK) regulates cell differentiation and gene expression in vertebrates’ biomineralizing cells, yet, little is known on ROCK’s role in invertebrates’ biomineralization. Here we reveal that ROCK controls the formation, growth and morphology of the calcite spicules in the sea urchin larva. ROCK expression is elevated in the sea urchin skeletogenic cells downstream of the Vascular Endothelial Growth Factor (VEGF) signaling. ROCK inhibition leads to skeletal loss and disrupts skeletogenic gene expression. ROCK inhibition after spicule formation reduces spicule elongation rate and induces ectopic spicule branching. Similar skeletogenic phenotypes are observed when ROCK is inhibited in a skeletogenic cell culture, indicating that these phenotypes are due to ROCK activity specifically in the skeletogenic cells. Reduced skeletal growth and enhanced branching are also observed under direct perturbations of the actomyosin network. We propose that ROCK and the actomyosin machinery were employed independently, downstream of distinct GRNs, to regulate biomineral growth and morphology in Eukaryotes.

## Introduction

Biomineralization is the process in which organisms from the five kingdoms of life, use minerals to produce shells, skeletons and teeth that protect and support them (1–4). Recent studies suggest that biomineralization evolved by the phylum specific co-option of ancestral gene regulatory networks (GRNs) that drove the construction of distinct organic scaffolds (4–9) and by the evolution of specialized sets of biomineralization proteins (3, 7, 10–13). This explanation is in line with the dissimilar GRNs and biomineralization proteins that drive biomineralization in different phyla (5, 12, 14). Yet, there are common cellular processes required for mineralization in highly diverse phyla (4), suggesting that distinct upstream GRNs might have recruited common cellular and molecular mechanisms to drive biomineralization.

The actomyosin network was shown to play a role in biomineralization in various Eukaryote models, from unicellular organisms to vertebrates’ bones and teeth (15–22). There is a tight association between actin filaments and the biomineralization compartment in diatoms (19) and a highly dynamic actin organization that forms the biomineralization compartment in foraminifera (17). Perturbations of actin polymerization result with severely deformed shells and inhibition of shell shedding in coccolithophores (15, 16) and diatoms (18). In vertebrates, Rho GTPases and Rho-associated coiled-coil kinase (ROCK), regulate chondrocytes, osteoblasts and odontoblasts differentiation and affect genes expression in these biomineralizing cells (20–22). However, the roles of actomyosin remodeling in controlling mineral deposition and shaping biomineral morphology, are still unclear.

Sea urchin larval skeletogenesis provides an attractive model to study the role of the actomyosin network in biomineralization. Sea urchin larval skeletons are made of two frameworks of interconnected calcite rods, termed “spicules” that are generated by the skeletogenic cells (6, 23). To make the spicules, the skeletogenic cells form a ring with two lateral skeletogenic clusters and fuse through their filopodia forming a pseudopodia cable that links them into a syncytium (24, 25). The mineral is concentrated in the form of amorphous calcium carbonate (ACC) inside intracellular vesicles (26, 27). The vesicles are secreted into the biomineralization compartment generated in the skeletogenic cell clusters forming two triradiate spicules (Fig. 8A, (26, 28)). The tubular biomineralization compartment, also called the spicule cavity, elongates within the pseudopodia cable by localized mineral deposition at the tip of each rod (29).

The GRN that controls sea urchin skeletogenesis is known in great detail and is highly similar to the GRN that controls vertebrates’ vascularization, suggesting a common evolutionary origin of these two tubulogenesis programs (6, 23, 30). As the spicules elongate, the expression of key regulatory and biomineralization related genes becomes restricted to the skeletogenic cells proximal to the growing tips, possibly to promote mineral deposition at these sites (31–33). Localized gene expression is regulated by signaling cues such as the Vascular Endothelial Growth Factor (VEGF) signaling (30–32, 34). However, how the skeletogenic GRN drives spicule formation and localized mineral deposition at the tips of the rods is poorly understood.

Previous works suggest that the actomyosin network is essential for sea urchin skeletal growth. Calcium bearing vesicles perform an active diffusion motion in the skeletogenic cells with a diffusion length that inversely correlates with the strength and activity of the actomyosin network (35). Actin filaments are formed around the spicule (35, 36) and F-actin signal is enriched at the tips of the growing skeletal rods in skeletogenic cell culture (37). Genetic and pharmacological perturbations of the GTPase, CDC42, prevent the formation of filopodia in the skeletogenic cells and inhibit spicule formation and elongation (38). Pharmacological perturbations of ROCK prevent spicule formation (39) and genetic perturbations of Rhogap24l/2 result with ectopic spicule splitting (6). Despite these characterizations, little is known on the role of the actomyosin machinery in regulating sea urchin biomineral growth and morphology.

Here we study the role of ROCK and the actomyosin network in the sea urchin *Paracentrotus lividus (P. lividus)*. Our findings reveal the critical role of ROCK and the actomyosin network in multiple aspects of sea urchin biomineralization, suggesting a common use of these factors in Eukaryote biomineralization, downstream of distinct GRNs.

## Results

### ROCK is enriched in the skeletogenic cells depending on VEGF signaling

We sought to study the spatial expression of the ROCK protein and its regulation by VEGF signaling, a prominent regulator of sea urchin skeletogenesis (6, 30). The sequence of ROCK and especially, its functional domains are highly conserved between human and sea urchin (Fig. S1, (40–42)). According to RNA-seq data measured in *P. lividus*, ROCK is a maternal gene that degrades after the maternal to zygotic transition and picks up again after hatching (>14 hours post fertilization (hpf), Fig. S2A). We used a commercial antibody that recognizes human ROCK protein to test ROCK expression in *P. lividus* embryos. We first used western blot to detect ROCK expression in control and under VEGFR inhibition at the time of spicule formation and during skeletal elongation (Axitinib treatment, ∼22hpf, 27hpf and 33hpf, Fig. S2D). The antibody detected a ∼150kDa protein (Fig. S2D) that is the predicted size for *P. lividus* ROCK protein (153kDa). VEGFR inhibition marginally increased the overall level of ROCK at 22hpf, but did not affect it at 27hpf and 33hpf (Fig. S2E). Yet, this measurement was done on proteins extracted from whole embryos, of which the skeletogenic cells, where VEGFR is active, are less than 5% of the total cell mass (43). We therefore wanted to study the spatial expression of ROCK and specifically, its regulation in the skeletogenic cells.

We studied the spatial distribution of ROCK protein using ROCK antibody and the skeletal cell marker, 6a9, that marks the skeletogenic syncytium membrane, in control and under VEGFR inhibition ((44), Fig. 1). We quantified ROCK signal in the skeletogenic cells compared to the neighboring ectodermal cells in both conditions, at the three time points mentioned above. In the three time points, under VEGFR inhibition, the skeletogenic cells do not migrate to their proper positions in the anterlateral and post-oral chains, as previously reported (6, 30, 34). At 22hpf, ROCK expression in the skeletogenic cells is mildly enriched compared to the neighboring ectodermal cells and this enrichment is not observed under VEGFR inhibition (Fig. 1A-F). At 27hpf, ROCK enrichment in the skeletogenic cells increases, but similar enrichment is observed under VEGFR inhibition (Fig. 1G-L). At 33hpf, ROCK enrichment in the skeletogenic cells is most apparent (Fig. 1M-O) and depends on VEGF signaling (Fig. 1P-S). At both 27hpf and 33hpf in control embryos, ROCK seems localized near the skeletogenic cell membranes, which further supports its activation in these cells, since ROCK activation leads to its localization to the cell membranes in other systems (45). Overall, this data demonstrate that ROCK expression is elevated in the skeletogenic cells and this enrichment strengthens with skeletal elongation and depends on VEGF signaling.

**Figure 1.**
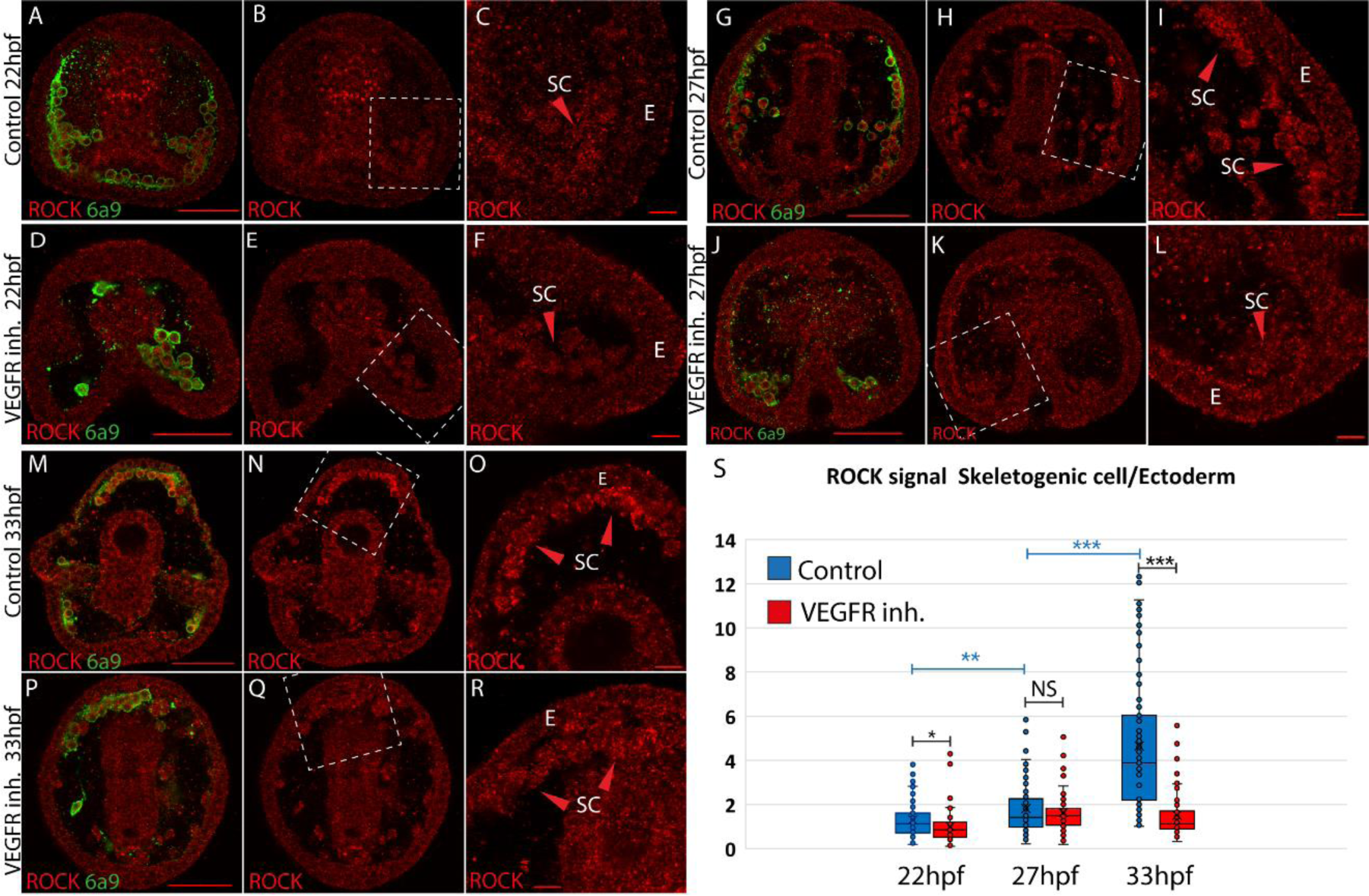
ROCK enrichment in the skeletogenic cells increases with time and depends on VEGF signaling. (A-R) ROCK immunostaining at different times points in control and under VEGFR inhibition (150 nM Axitinib). In each condition and time point, the left image shows ROCK immunostaining and the skeletogenic cell marker, 6a9 (A, D, G, J, M and P). The middle image shows ROCK immunostaining alone in the whole embryo (B, E, H, K, N and Q). The right image shows the enlargement of the white rectangle region of the middle image (C, F, I, L, O, R). Scale bar in whole embryo is 50µm and in enlargements is 10µm. E – ectoderm, SC – skeletogenic cells. (S) quantification of the ratio between ROCK signal/area in the skeletogenic cells compared to the ectodermal cells (see methods for details). Each box plot shows the average marked in x, the median, the first and the third quartiles (edges of boxes) and all experimental measurement (dots). Experiments were performed in three independent biological replicates and in each condition at least 33 embryos were measured. Statistical significance was measured using paired 2-tailed t-test where, * indicates P<0.05, ** indicates P<0.005, and *** indicates P<0.0005.

### ROCK activity in the skeletogenic cells controls spicule initiation, growth and morphology

Next, we studied the role of ROCK in sea urchin skeletogenesis using genetic perturbations. We downregulated the expression of ROCK by the injection of two different translation morpholino anti-sense oligonucleotides (MASOs). ROCK MASO-1 matches the start of translation region, ROCK MASO-2 matches the 5’ region upstream the start of translation, and random MASO was used as a control. Embryos injected with either ROCK MASO-1 or MASO-2 show reduced ROCK signal at 33hpf, supporting the downregulation of the ROCK protein using these MASO’s (Fig. S3B, F, J, N, R, V).

Embryos injected with ROCK MASO-1 or MASO-2 show strongly reduced skeletons at two days post fertilization (2dpf, Fig. 2C, D) and embryos injected with ROCK MASO-2 show an additional phenotype of severely branched skeleton at this time (Fig. 2B). The additional branching phenotype and the larger percentage of affected embryos in ROCK MASO-2 are probably due to its higher efficiency resulting from its more 5’ location relative to translation start codon (Fig. 2E). Thus, the genetic perturbations of ROCK expression indicate that sea urchin ROCK is important for skeletal elongation and normal branching pattern.

**Figure 2.**
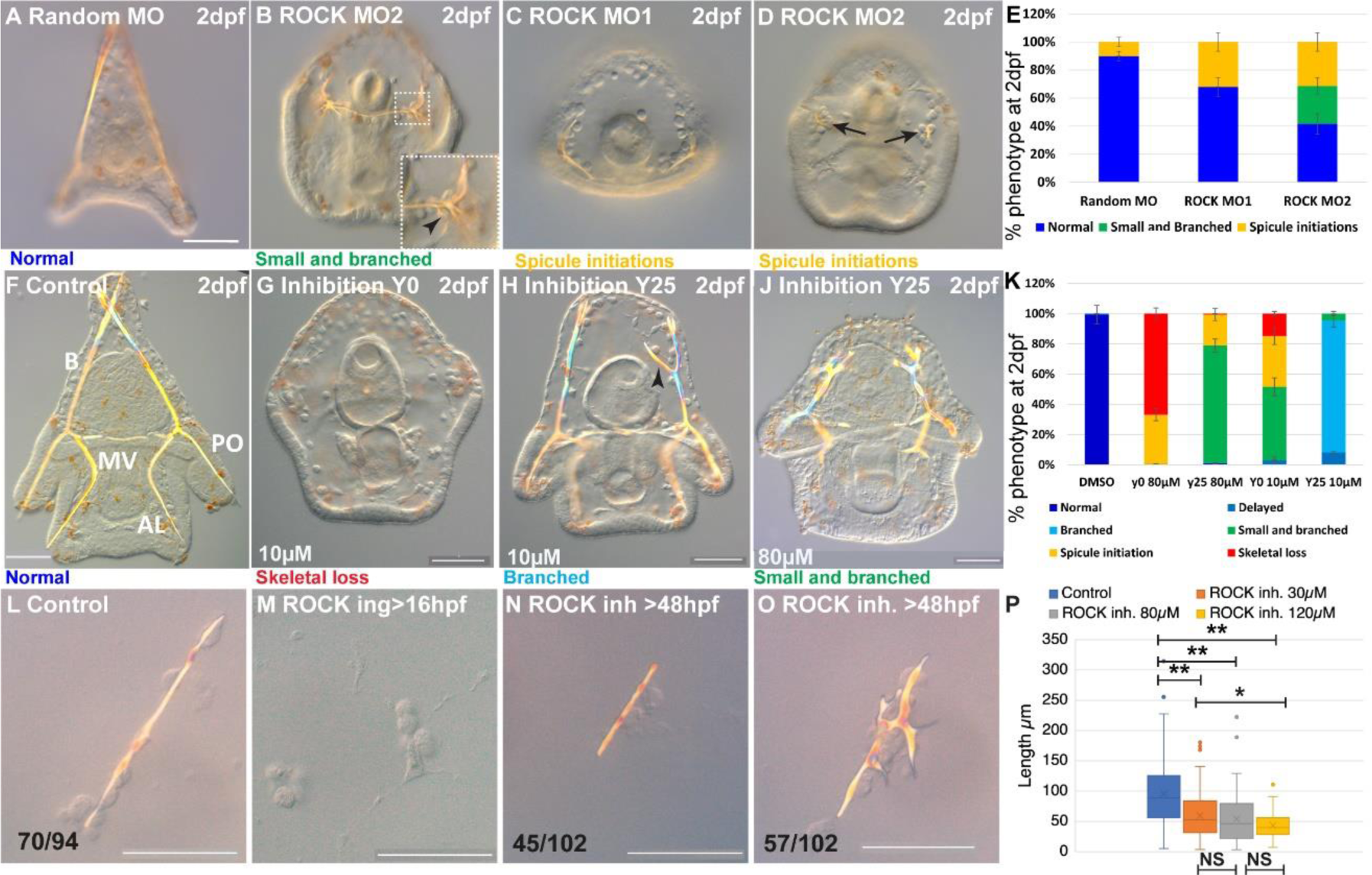
ROCK activity is essential for spicule formation, normal elongation and branching in whole embryos and in skeletogenic cultures. (A-E) genetic perturbation of ROCK translation using two different MASOs observed at 2dpf. (A) control embryo injected with Random MASO. (B) embryo injected with ROCK MO-2 shows ectopic spicule branching. (C, D) embryos injected with ROCK MO-1 or MO-2 show spicule initiations. (E) summary of MASO injection phenotypes based on 4-6 independent biological replicates. (F-K) pharmacological perturbations of ROCK activity using 10µM and 80µM of the inhibitor Y27632 observed at 2dpf. (F) representative control embryo with normal skeletal rods, B, body; AL, anterolateral; PO, post-oral and MV, midventral. (G) complete skeletal loss in embryo treated continuously with 10µM ROCK inhibitor. (J) reduced skeletal growth and enhanced ectopic branching in embryo where 10µM ROCK inhibitor was added at 25hpf. (J) small spicules with enhanced ectopic branching in embryo where 80µM ROCK inhibitor was added at 25hpf. (K) summary of perturbation phenotypes based on 3-8 biological replicates for each treatment. See additional concentrations, phenotypes and summary in Fig. S4 and Table S1. (L-O) representative spicules from skeletogenic cell cultures in control and under 30µM Y27632 at 72hpf. (L) linear spicule in control culture, (M) Y27632 addition at 16hpf, before spicule initiation, completely blocks spiculogenesis. (N, O) Y27632 addition after spicule initiation, at 48hpf, reduces spicule elongation (N), and enhances branching (O). (P) quantification of spicule length in control and ROCK inhibition (>48hpf) at 72hpf. *P < 0.05, **P < 0.001, Kruskal-Wallis non-parametric test. Results are based on three biological repeats for each treatment, except from 120µm that was done in two biological repeats. Scale bars are 50µm. In L, N and O, the numbers at the bottom indicate the number of spicules that show this phenotypes (left) over all observed spicules (right).

To elucidate the dependence of ROCK phenotypes on ROCK activity level and identify its effect on different stages of skeletogenesis, we tested the skeletogenic phenotypes of different concentrations of ROCK inhibitor, Y27632, applied at different time points (Fig. S4A). Y27632 binds to the ATP site of ROCK’s kinase domain and prevents its activity (46). Y27632 affinity to ROCK is 100 times higher than its affinity to other kinases, such as PKA and PKC (47). The amino-acids to which Y27632 binds, are conserved in the sea urchin ROCK protein, supporting its specific inhibition of this protein (Fig. S1C). Y27632 had been used in cell cultures and live embryos from vertebrates to *Drosophila,* in concentrations between 10-100µM (47–49). In the sea urchin embryo a concertation of 75µM of Y27632 was reported to completely block skeleton formation (39) and a concentration of 100µM was shown to delay gut invagination (50). Therefore, we tested the range of 10-80µM Y27632 applied before or after spicule initiation (Fig. S4A).

Continuous ROCK inhibition beginning at egg fertilization, resulted with significant skeletogenic phenotypes that are dose dependent (Fig. 2, Fig. S4). Continuous ROCK inhibition using 80µM Y27632, did not affect skeletogenic cell migration but eliminated skeletal formation at 27hpf (Fig. S4B, C). At 2dpf, complete skeletal loss is detected under continuous ROCK inhibition in all concentrations, ranging from 15% of the embryos exposed to 10µM to 67% of the embryos exposed to 80µM Y27632 (Fig. 2G, K, in agreement with (39)). The rest of the embryos exposed to continuous ROCK inhibition show either spicule initiations or small spicules with enhanced ectopic branching, with percentage depending on Y27632 concertation (Fig. 2K, S4K). Except from the skeletogenic phenotypes, the overall embryonic development of embryos exposed to Y27632 in all concentrations seems normal (Fig. 2G, J). Similar results are observed when adding the inhibitor at 20hpf, with distribution of phenotypes that depends on the concentration (Fig. S4I, K). Importantly, skeletal loss, spicule initiation and ectopic branching were not observed under PKC or PKA inhibition, that resulted with much milder skeletogenic phenotypes (PKC) or no skeletogenic phenotype (PKA, (51, 52)), supporting the selective inhibition of ROCK by Y27632. Altogether, continuous ROCK inhibition results with severe skeletogenic phenotype ranging from complete skeletal loss to small spicule with ectopic branching, with ratios that depend on the inhibitor concentration.

The addition of the inhibitor at 25hpf, after spicule initiation, results in a majority of embryos showing ectopic spicule branching (Fig. 2H-K, S4F-H, K). The reduction of skeletal growth rate and ectopic branching can be observed a few hours after the addition of ROCK inhibitor (Fig. S5, 30µM). Washing the inhibitor at the highest concertation after 25hpf results in partial recovery of skeletogenesis, with normal or mildly delayed skeletons, indicating that ROCK inhibition is reversable (Fig. S4E, K).

Overall, the genetic and pharmacological perturbations of ROCK result with sever reduction of skeletal growth and enhanced skeletal branching, but only continuous ROCK inhibition leads to complete skeletal loss (Fig. 2G, K). Immunostaining of fertilized eggs clearly shows that ROCK protein is maternal, in agreement with our RNA-seq data (Fig. S2A-C). The injected MASO’s cannot interfere with the maternal ROCK protein whereas the inhibitor affects the activity of both maternal and zygotic ROCK, which could underlie the absence of skeletal loss in the genetic perturbations. Indeed, ROCK signal is only moderately reduced in ROCK MASO injected embryos at 33hpf (Fig. S3F, J, N), demonstrating the partial penetration of the MASO. Thus, our genetic and pharmacological perturbations reconcile, and indicate that ROCK activity is necessary for spicule formation, skeletal elongation and normal branching pattern.

To test if ROCK skeletogenic phenotypes are due to its activity specifically in the skeletogenic cells, we inhibited ROCK in a culture of isolated skeletogenic cells (Fig. 2L-P). The addition of ROCK inhibitor to the skeletogenic cell culture at 16hpf, before the spicules form, completely abolished spicule formation (Fig. 2M). The addition of the inhibitor at 48hpf, after spicule formation, resulted with significantly shorter spicules (Fig. 2L, N) and increased branching compared to control spicules (Fig. 2O). The reduction of the spicule length becomes more significant with increasing concentration of the ROCK inhibitor (Fig. 2P). Notably, skeletal loss and ectopic spicule branching were not observed in PKA or PKC inhibition in skeletogenic cell cultures (52), further supporting the selective inhibition of ROCK by Y27632. ROCK skeletogenic phenotypes in the isolated skeletogenic cell culture are similar to its phenotypes in whole embryos, verifying that ROCK activity in the skeletogenic cells is essential for spicule formation, normal elongation and prevention of branching.

### ROCK inhibition reduces spicule volume, surface area and total length, but not thickness (SR-µCT)

To quantify the effect of ROCK inhibition on mineral deposition and skeletal shape we used synchrotron radiation micro-computed tomography (SR-µCT) that enables three-dimensional visualization and quantification of spicule morphology at 2 and 3dpf (Figs. 3, S6, 40µM Y27632, see methods for details). The aberrant morphology of ROCK inhibited spicules is visible at both developmental time points (Fig. 3A). The ectopic branching occurs both at the tips where tip-splitting is detected (arrowheads, Fig. 3A), as well as in the back, where small spicule extensions, or mineral ‘dripping’ are observed (arrow at 48hpf, Fig. 3A). This demonstrates that the regulation of mineral deposition is perturbed both at the tip and in the back of the spicules under ROCK inhibition.

**Figure 3.**
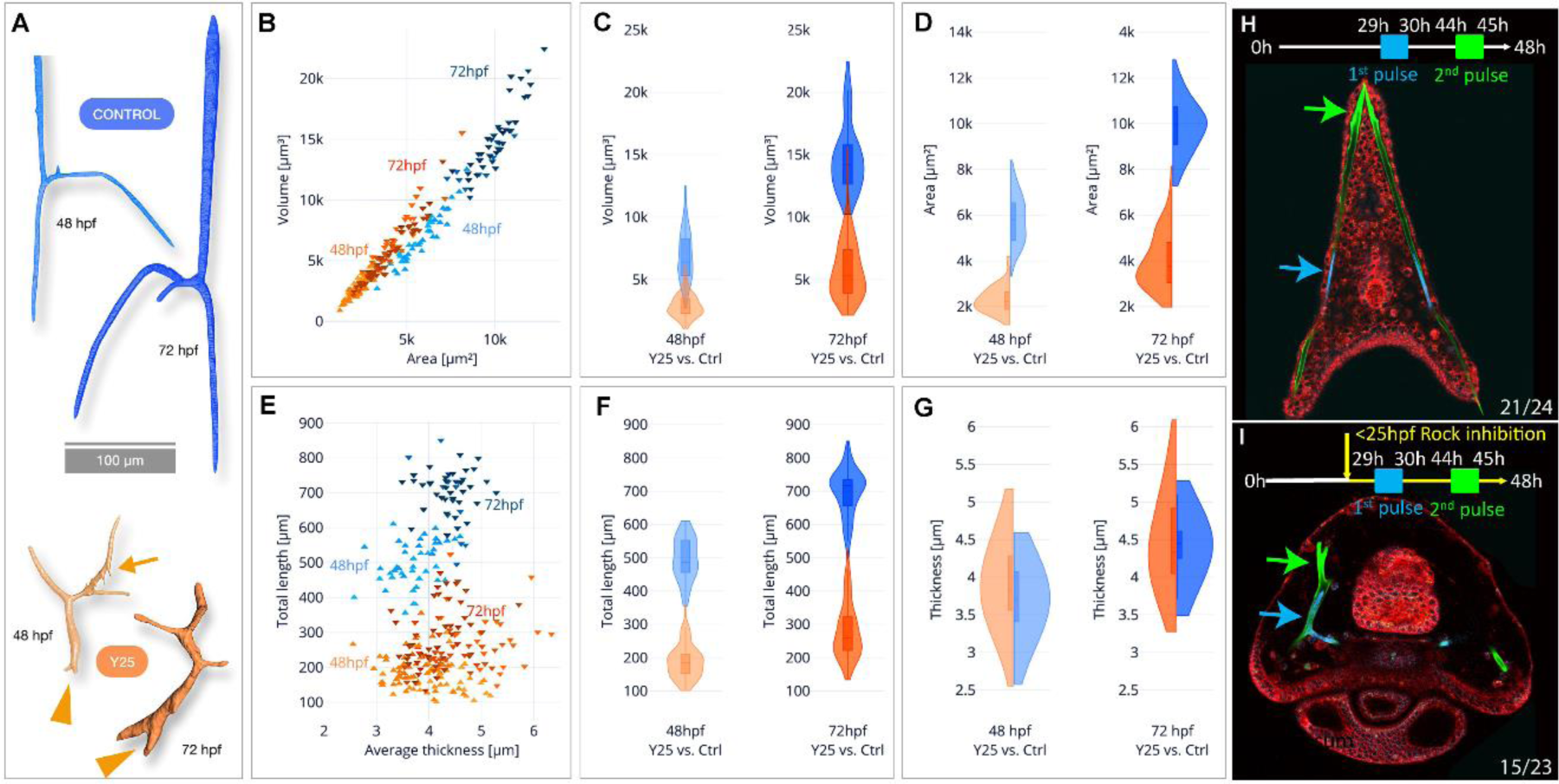
SR-µCT studies of ROCK inhibited spicuels show reduction in skeletal volume, surface area and total length, but not thickness and 2-pulse calcein shows loss of tip-dominance. (A) Exemplary 3D-renderings of control spicules (top, blue) and spicules where 40µM of ROCK inhibitor was added at 25hpf (bottom, orange), dissected at 48hpf and 72hpf. Arrowheads point to tip splitting and arrow at 48hpf point to spicule dripping at the back. (B) Spicule volume vs. area for control and ROCK-inhibited spicules at 48 and 72 hpf. Each data point represents a single spicule. (C-D) Frequency distributions for volume and surface area of control and ROCK-inhibited spicules at 48 hpf and 72 hpf (left and right violin plots, respectively). (E) Spicules’ total branch length and average thickness for control and ROCK-inhibited spicules at 48 and 72 hpf. (F-G) Frequency distributions for spicule lengths and thickness of control and ROCK-inhibited spicules dissected at 48hpf and 72hpf. (H, I) Calcein two-pulses experiment. Embryos were exposed to blue calcein at 29-30hpf, to green calcein at 44-45hpf and stained with FM4-64 membrane marker (red) prior to image acquisition at 48hpf. (H) Control embryo, (G) Embryo where 30µM of Y27632 was added at 25hpf. The experiments were done in three biological replicates and the numbers at the bottom indicate the number of embryos that show this phenotype out of all embryos scored.

When compared to control spicules, ROCK inhibition leads to a ∼2.5-fold reduction of the spicule volume and surface area in both 2 and 3dpf (Fig. 3B-D, see Table S2 for measured values and standard deviations and S3 for statistical analyses). The average rate of spicule growth between 48hpf and 72hpf in control embryos is 325.5µm^3^/hour, and reduces to 119.8µm^3^/hour under ROCK inhibition. Hence, the rate of mineral deposition is significantly reduced under ROCK inhibition.

To understand the specific effect of ROCK inhibition on spicule elongation vs. thickening, we compared the total length and the mean thickness of the spicules between control and ROCK inhibition (Fig. 3E-G). The total length of control spicules was on average about 2.5-times longer than the length under ROCK inhibition in both 2dpf and 3dpf (Fig. 3E, F, tables S2 and S3). In contrast to its effect on the spicule length, ROCK inhibition caused a minor increase of the spicule thickness at 2dpf and at 3dpf the thickness was not affected by the inhibition (Fig. 3E-G, Tables S2, S3). These SR-µCT measurements indicate that ROCK activity plays a crucial role in controlling the rate of mineral deposition at the tips of the rods and spicule elongation, and is not affecting spicule thickening.

### ROCK activity does not control mineral uptake but is required for “tip-dominance”

To test the effect of ROCK inhibition on mineral intake, we used the fluorescent chromophore, calcein that binds to divalent ions including Ca^+2^ and is widely used to track the calcium pathway in biomineralization (6, 28, 33, 35). We measured the number of calcein stained vesicles per area as well as the number of pixels of calcein signal per area in the skeletogenic cells, in control and under ROCK inhibition at 1dpf and 2dpf (Fig. S7A-E). Continuous ROCK inhibition does not change the number of calcium bearing vesicles at 1dpf, but the number of calcein stained pixels significantly increases in this condition, possibly indicating that the vesicle volume is larger (Fig. S7F, G). At 2dpf, the number of calcein stained pixels is similar in control and in continuous ROCK inhibition, suggesting that the overall calcium content does not change in this condition (Fig. S7H, I). The number of calcein stained vesicles is however decreased under ROCK inhibition, possibly indicating that there are fewer vesicles with larger volume. Addition of ROCK inhibitor at 25hpf affects neither the number of calcein stained vesicles nor the number of calcein stained pixels in the skeletogenic cells. Together, these measurements show that ROCK is not required for the uptake of calcium into the cells. Therefore, the skeletal phenotypes occurring under ROCK inhibition are not related to a decrease in calcium uptake or shortage in intracellular calcium, but are due to defects in mineral processing within the cells and in mineral deposition.

To monitor the effect of ROCK inhibition on mineral deposition rate and distribution, we applied two pulses of different calcein dyes (53): blue calcein was added at 29hpf and washed after one hour, followed by green calcein added at 44hpf and washed after one hour. Hence, blue calcein marks the initial triradiate region and green calcein marks the edges of the spicules that were recently deposited (blue and green arrows in Fig. 3H, I). The dye FM4-64 was used to mark the cell membranes. Under ROCK inhibition, the green labeled region is much closer to the blue labeled region compared to control, due to the reduction in skeletal elongation rate, in agreement with our SR-µCT measurements. However, while in control embryos each body rod has a single tip stained in green calcein, under ROCK inhibition one of the body rods has two tips and both are stained in green. This indicates that mineral is being deposited in both edges, and the mechanism that prevents the growth of multiple tips in each rod, enabling “tip-dominance”, (in analogy to plant stem apical dominance), is disrupted under ROCK inhibition.

### The activity of the actomyosin network is essential for normal spicule elongation and inhibition of ectopic branching

In other systems, major roles of ROCK are to control F-actin polymerization and MyoII activation (54–58), hence we wanted to directly test the role of these actomyosin components in sea urchin skeletogenesis. To directly inhibit F-actin polymerization we used Latrunculin-A (Lat-A), that prevents the polymerization of actin monomers (59) and was shown to be effective in sea urchin embryos (60). To directly inhibit actomyosin contractility we used Blebbistatin (Blebb), that interferes with MyoII ATPase activity, prevents actin-myosin interactions (61) and was shown to be effective in sea urchin embryos (62). To prevent interference with the early cell divisions where the actomyosin network plays a critical role (63–65), we added the inhibitors, individually or together, before or after spicule formation (20hpf and 25hpf, see methods for details). We tested the resulting phenotypes at 2dpf (Fig. 4A-G).

**Figure 4.**
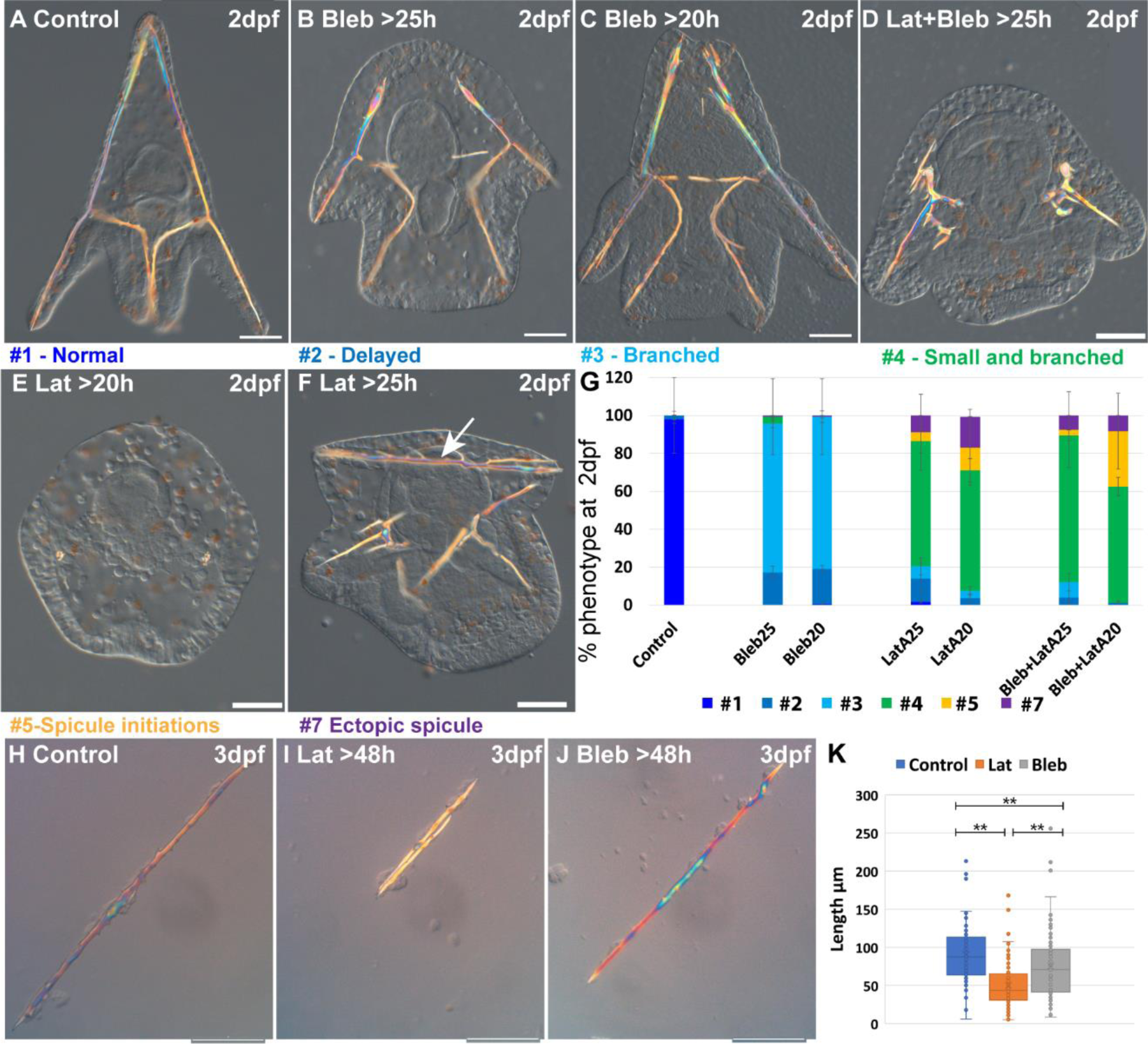
Actin polymerization and myosin activation affect skeletal growth and branching. (A-F) representative embryos showing the effect of actomyosin perturbations at 2df. (A) control embryo (B) embryo treated with 2μM Blebbistatin >25hpf , (C) embryo treated with 2μM Blebbistatin >20hpf, (D) embryo treated with 2nM Latrunculin-A and 1.5μM Blebbistatin >25hpf, (E) embryo treated with 2nM Latrunculin-A >20hpf, (F) embryo treated with 2nM Latrunculin-A >25hpf, arrow pointing to the additional spicule rod. (G) Statistics of Latrunculin-A and Blebbistatin phenotypes, color code of phenotype is indicated in the representative images. Error bars indicate standard deviation. All treatments were conducted in at least three biological replicates and exact number of replicates and scored embryos are provided in table S4. (H-J) representative spicules recorded at 72hpf from (H) control skeletogenic cell culture, (I) skeletogenic cell culture were 2nM Latrunculin-A was added at 48hpf and (J) skeletogenic cell culture were 2μM Blebbistatin was added at 48hpf. (K) Quantification of spicule length in the different treatments at 72hpf **P < 0.001, Kruskal-Wallis non-parametric test. Results are based on three biological repeats for each treatment. Scale bars are 50μm.

Our results indicate that F-actin polymerization and to a lesser extent, actomyosin contractility, are essential for normal skeletal growth and inhibition of ectopic branching. The majority of the embryos where MyoII activation was inhibited show ectopic branching mostly at the tips of the rods, and a minor delay in skeletal growth (Fig. 4B, C, G). These phenotypes are quite similar to those observe in the addition of 10µM of ROCK inhibitor at 25hpf (Fig. 2H, K). The majority of the embryos where F-actin formation was inhibited show significant reduction of skeletal growth and severe ectopic skeletal branching (Fig. 4E, F and similar phenotypes to Fig. 4D, statistics in Fig. 4G). These phenotypes are quite similar to those observed in 10µM continuous inhibition of ROCK and the addition of 30µM of ROCK inhibitor at 20hpf (Fig. 2H, K). In some embryos, additional spicules form (Fig. 4F, G). The co-inhibition of F-actin polymerization and MyoII activity results in skeletogenic phenotypes that resemble, but are slightly stronger, than those of the inhibition of F-actin alone (Fig. 4D, G). Thus, the phenotypes resulting from the inhibition of MyoII activation resemble those of the late inhibition of ROCK at low inhibitor concertation. The phenotypes resulting from inhibition of actin polymerization resemble those of the continuous inhibition of ROCK in low inhibitor concertation and the late inhibition of ROCK at higher inhibitor concentrations, yet skeletal loss is only observed under ROCK continuous inhibition.

To test whether actin polymerization and myosin activity specifically in the skeletogenic cells underlies the observed skeletogenic phenotypes, we inhibit both in an isolated skeletogenic cell culture (Fig. 4H-K). We added the inhibitors after the spicule formed (>48hpf) and observed the phenotypes at 3dpf. Both inhibitors reduced skeletal growth, with Lantruculin-A having a significantly stronger effect than Blebbistatin (Fig. 4K), in agreement with the stronger skeletogenic phenotypes of Lantruculin-A in whole embryos. However, differently than in whole embryos, skeletal branching in both Lantruculin-A and Blebbistatin is not affected and is similar to the observed branching in control cultures. The absence of the branching phenotype in the skeletogenic cell culture could be due to the increased rigidity of the substrate that could compensate for the reduce actomyosin activity. Another option is that the branching phenotype in whole embryos is due to the reduction of the activity of the actomyosin network in non-skeletogenic cells. Nevertheless, these results indicate that normal skeletal growth depends primarily on F-actin polymerization and to a lesser extent, on myosin contractility in the skeletogenic cells.

### ROCK activity is required for F-actin organization around the forming spicule

The similarity between the skeletogenic phenotypes under ROCK inhibition and the direct perturbations of the actomyosin network led us to test the effect of ROCK inhibition on F-actin organization and MyoII activity. To accomplish this we used phalloidin, a fluorophore that binds F-actin specifically, an antibody that detects phosphorylated MyoII (MyoIIP, (35)) and the skeletogenic marker, 6a9 (Fig. 5). In control embryos, F-actin is detected around the tri-radiate calcite spicules at 27hpf (green arrow in Fig. 5B, in agreement with (35, 36)). The pseudopodia cable that connects the skeletogenic cells shows lower F-actin signal compared to the signal around the spicule (blue line in Fig. 5B, C, E blue arrows in Fig. 5G, H, J). MyoIIP signal is not enriched in the skeletogenic cells nor in the spicule (Fig. 5D, I). Under continuous ROCK inhibition, the pseudopodia that connects the skeletogenic cells still forms (Fig. 5R) and enhanced F-actin signal is still observed in the lateral skeletogenic cell clusters (green arrow in Fig. 5L); but unlike control embryos, the spicule cavity is not formed and F-actin is not organized around it (Fig. 5L). MyoIIP signal seems unchanged by ROCK inhibition at this time (Fig. 5N, S), in agreement with the weak and late phenotype of MyoII inhibition (Fig. 4). Thus, the pseudopodia cable forms but F-actin organization around the spicules does not occur under continuous ROCK inhibition, however it is not clear if the effect on F-actin organization is direct or due to the absence of spicule in this condition.

**Figure 5.**
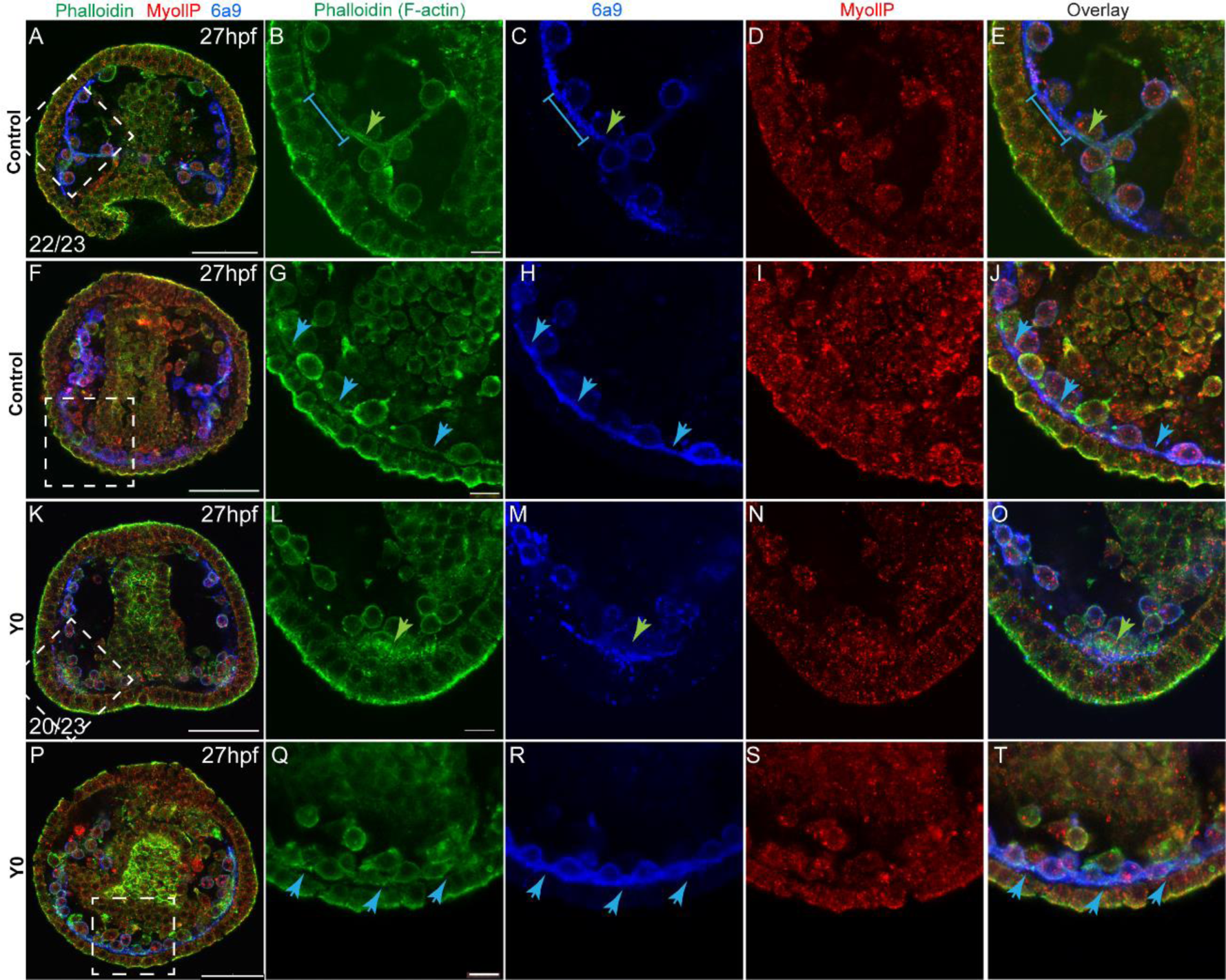
ROCK inhibition effect on F-actin organization and MyoII activity *at 27hpf*. Representative images showing normal embryo (A-J) and embryos treated with ROCK inhibitor from fertilization (K-T, 80µM), at 27hpf. Phalloidin (green) was used to stain F-actin, MyollP (red) was used to stain phosphorylated myosinII and 6a9 (blue) was used to mark the skeletogenic cells. Right panels show enlargements of the rectangle sections marked in the left most panels. Green arrows point to regions that show enriched F-actin signal, blue arrows point to the pseudopodia cable. The blue line in B, C and E marks a region of the pseudopodia cable that is stained by 6a9 but has low F-actin signal. The experiments were done in 3 biological replicates, the numbers at the bottom left of (A, K) indicate the number of embryos that show this phenotype out of all embryos scored. Scale bar in A, F, K and P is 50μm, and in B, I, N and S is 10μm.

### F-actin is enriched at the tips of the elongating spicules independently of ROCK activity

To assess the role of ROCK in actomyosin organization during skeletal elongation we compared F-actin and myoIIP signals between control embryos and embryos where ROCK inhibitor was added at 25hpf (Fig. 6). At the prism stage, we did not detect a clear difference in MyoIIP signal in the skeletogenic cells, between control and ROCK inhibited embryos (Fig. 6D, I). At this time, F-actin is enriched at the tips of the spicules in both control and ROCK inhibited embryos (white arrowheads in Figs. 6B, G) as well as in ROCK morphants (green arrowheads in Fig. S3K, W). In both control and ROCK inhibited embryos, the F-actin signal was markedly higher in the regions of the pseudopodia cable where the spicule cavity had formed, compared to the regions where the spicule cavity had not yet formed (blue arrows in Fig. 6B-E, G-J). We quantified the F-actin signal at the tips compared to the back in control and ROCK inhibited embryos (Fig. 6K, L, see methods for details). The phalloidin signal is on average 3-fold stronger at the tips of the spicules compared to the back, in both control and ROCK inhibition, indicating that F-actin enrichment at the tips is significant and independent of ROCK activity.

**Figure 6.**
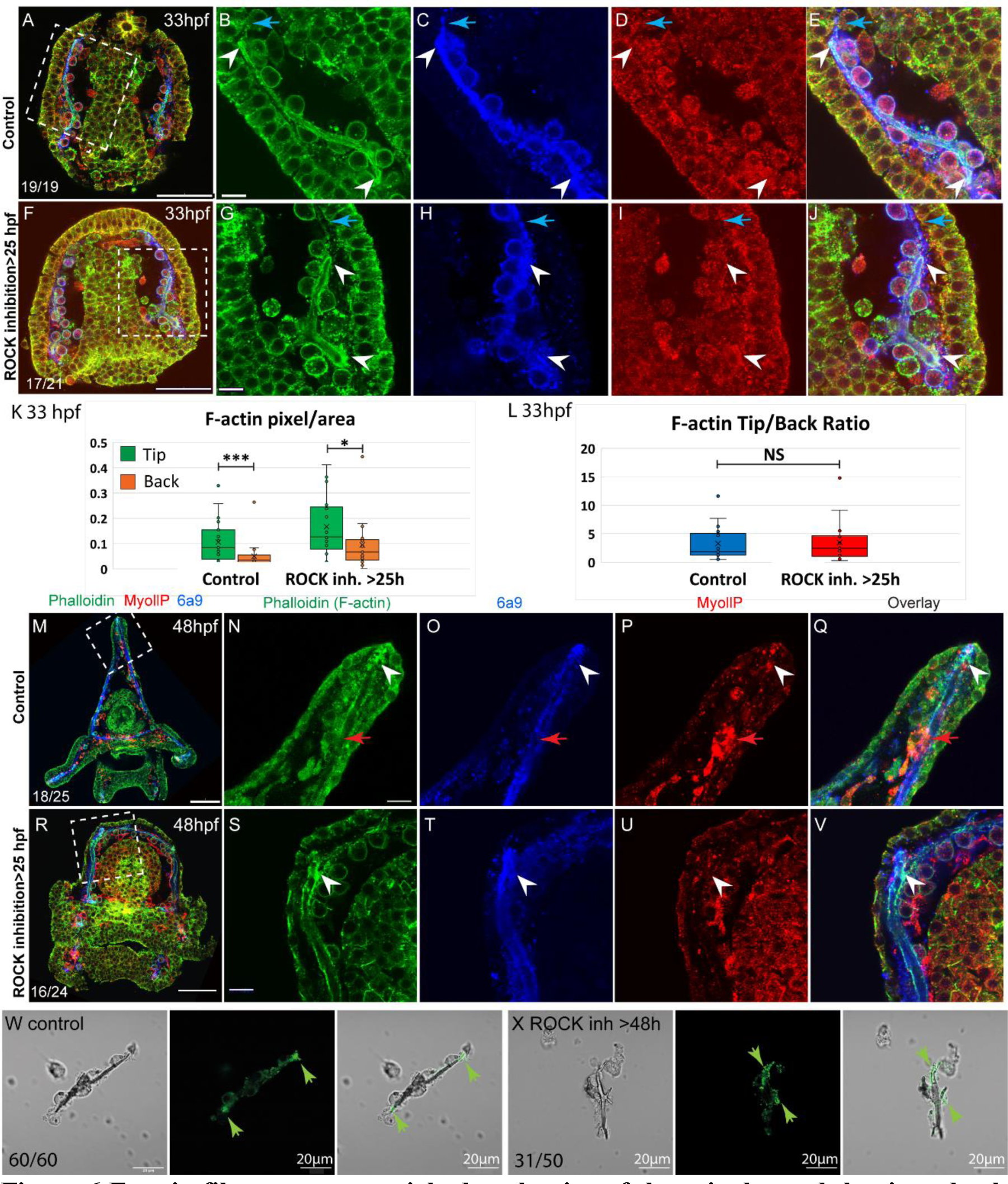
F-actin filaments are enriched at the tips of the spicules and the tip to back ratio is unaffected by ROCK inhibition. (A-J) representative images at 33hpf, showing normal embryo (A-E) and embryos treated with 30µM ROCK inhibitor from the gastrula stage (>25hpf, F-J). Embryos are stained with Phalloidin (green), MyollP antibody (red) and 6a9 (blue). (B-E, G-J) enlargement sections of the spicule area marked with rectangle in A and F. White arrowheads point to the enriched F-actin signal at the tips. Blue arrows point to the region of the pseudopodia cable that is not filled with the spicule cavity. (K, L) quantification of the tip to back F-actin signal (number of green pixels per area) at 33hpf in control embryos and ROCK inhibition >25hpf. Each box plot shows the average (x), median (middle line), the first and the third quartiles and all the points measured. Asterisks indicate statistical significance in paired t-test, where * is p<0.05 and *** is p< 0.0005, NS-not significant. (M-V) similar experiments to (A-J), at 48hpf, 40µM of ROCK inhibitor added at 25hpf. Red arrows point to non-skeletogenic cells enriched with MyoIIP. The experiments were repeated in 3 biological replicates and the numbers at the bottom left of (A, F, M, R) indicate the number of embryos that show this phenotype out of all embryos scored. Scale bar in A, F, M, R is 50µm and in B, G, N, S, is 10µm. (W, X) representative spicules out of three biological replicates from control skeletogenic cell culture (W, n=60), and skeletogenic cell treated with 30µM ROCK inhibitor added at 48hpf and recorded at 72hpf (X, n=52). Left panel is phase image, middle panel is phalloidin staining and right panel shows the overlay. Green arrows point to the enhanced F-actin signal at the tips. Scale bar is 20μm.

At the pluteus stage, F-actin is enriched at the tips of the spicules in both control and ROCK inhibited embryos (white arrowheads in Fig. 6N-Q, S-V). Some non-skeletogenic cells that are enriched with MyoIIP signal are detected at this time (red arrowheads in Fig. 6N-Q). Together these data demonstrate that F-actin coats the spicule cavity and is enriched at the tip of the rods, independently of ROCK activity.

We tested the effect of ROCK inhibition on F-actin in skeletogenic cell cultures. F-actin is detected at the tips, in both control cultures and cultures where ROCK was inhibited after spicule formation (Fig. 6W, X). Yet, under ROCK inhibition, branching is enhanced and F-actin is enriched at the splitting tips of the spicule rods (Fig, 6X). The enrichment of F-actin at the splitting tips of the spicule under ROCK inhibition in skeletogenic cell culture resembles the calcein staining at the two tips in the embryo in this condition (Fig. 3I). These observations further support the role of ROCK activity in regulating tip-dominance during sea urchin skeletogenesis.

### ROCK activity is essential for normal skeletogenic gene expression

The role of ROCK in regulating gene expression during vertebrates’ chondrocytes, osteoblasts and odontoblasts differentiation (20–22) intrigued us to study the effect of ROCK inhibition on skeletogenic gene expression in the sea urchin embryo. ROCK continuous inhibition significantly downregulates the expression of multiple skeletogenic genes, including the cytoskeleton remodeling gene, *rhogap24l/2*, the biomineralization genes, *caral7, scl26a5* and *SM30* at 27hpf and 2dpf (Fig. 7A, (6)). ROCK inhibition did not affect the expression level of VEGFR and ROCK itself at these times (Fig. 7A). Washing the inhibitor after 25hpf or adding the inhibitor at 25hpf had a weaker effect, but still led to downregulation of *caral7*, *SM30*, *angio1* and *notch1* (Fig. 7B). Thus, ROCK activity is required for the normal expression level of multiple skeletogenic genes during sea urchin skeletogenesis.

**Figure 7.**
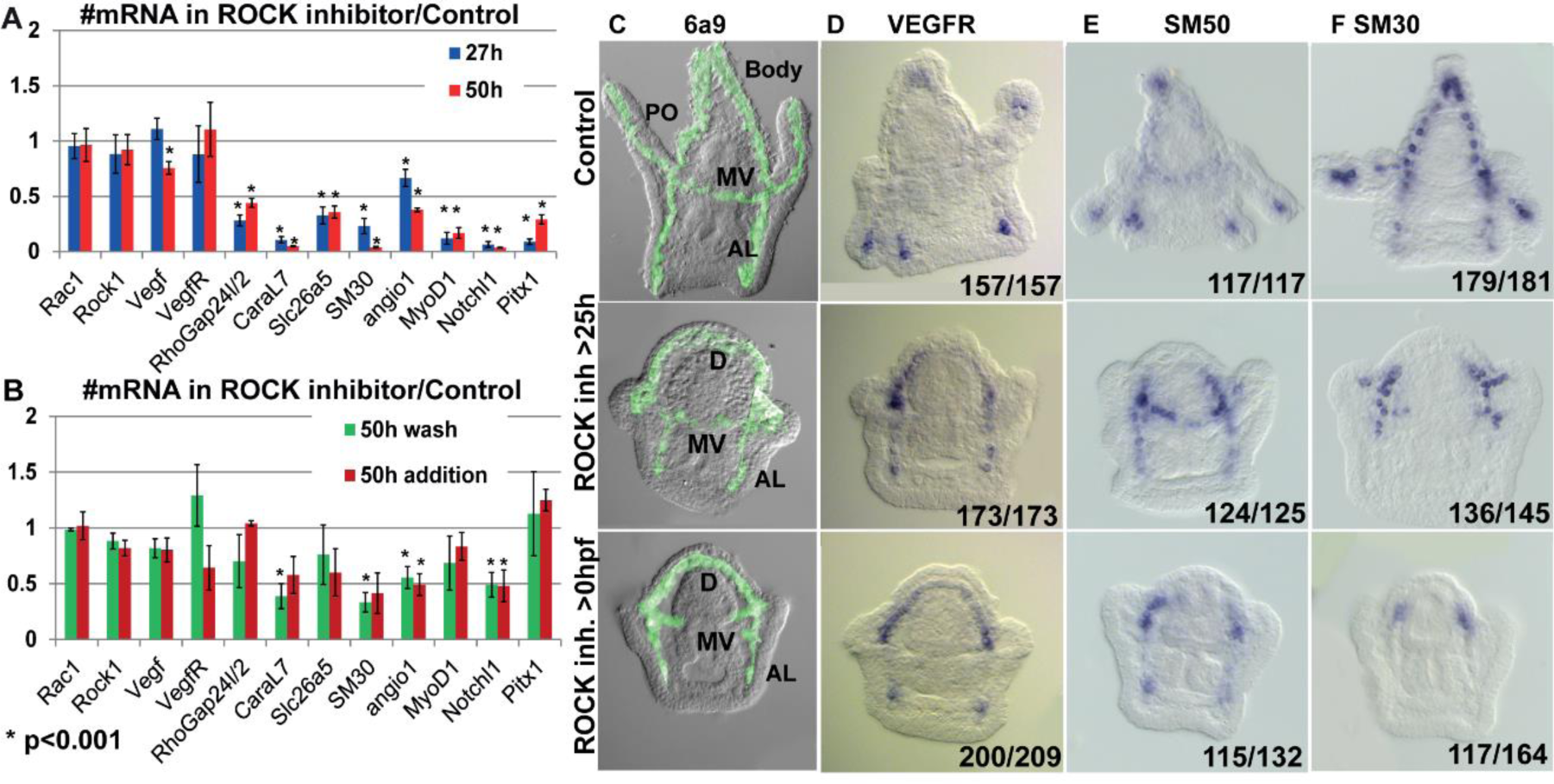
ROCK activity is essential for normal gene expression in the skeletogenic cells. (A, B) the effect of 80µM ROCK inhibition on gene expression. (A) continuous ROCK inhibition at 27hpf and 50 hpf (n=4). (B) addition of ROCK inhibitor at 25hpf and the wash of ROCK inhibitor at 25hpf, measured at 50hpf (n=3). Asterisks indicate p<0.001, one tailed z-test. Error bars show standard deviation. (C-F) Representative images of control embryo (top panels), embryos where ROCK inhibitor was added at 25hpf (middle panels), and embryos that were exposed to continuous ROCK inhibition (bottom panels), at the pluteus stage (∼48hpf). (C) skeletogenic cell marker, 6a9. MV, midventral; AL, anterolateral, and PO, Post-oral rods, D, dorsal skeletogenic chain. (D-F) spatial expression of skeletogenic genes. Gene names are indicated at the top of each panel. Numbers at the bottom of each image indicate the number of embryos that show this phenotype (left) out of all embryos scored (right), conducted in at least three independent biological replicates.

**Figure 8.**
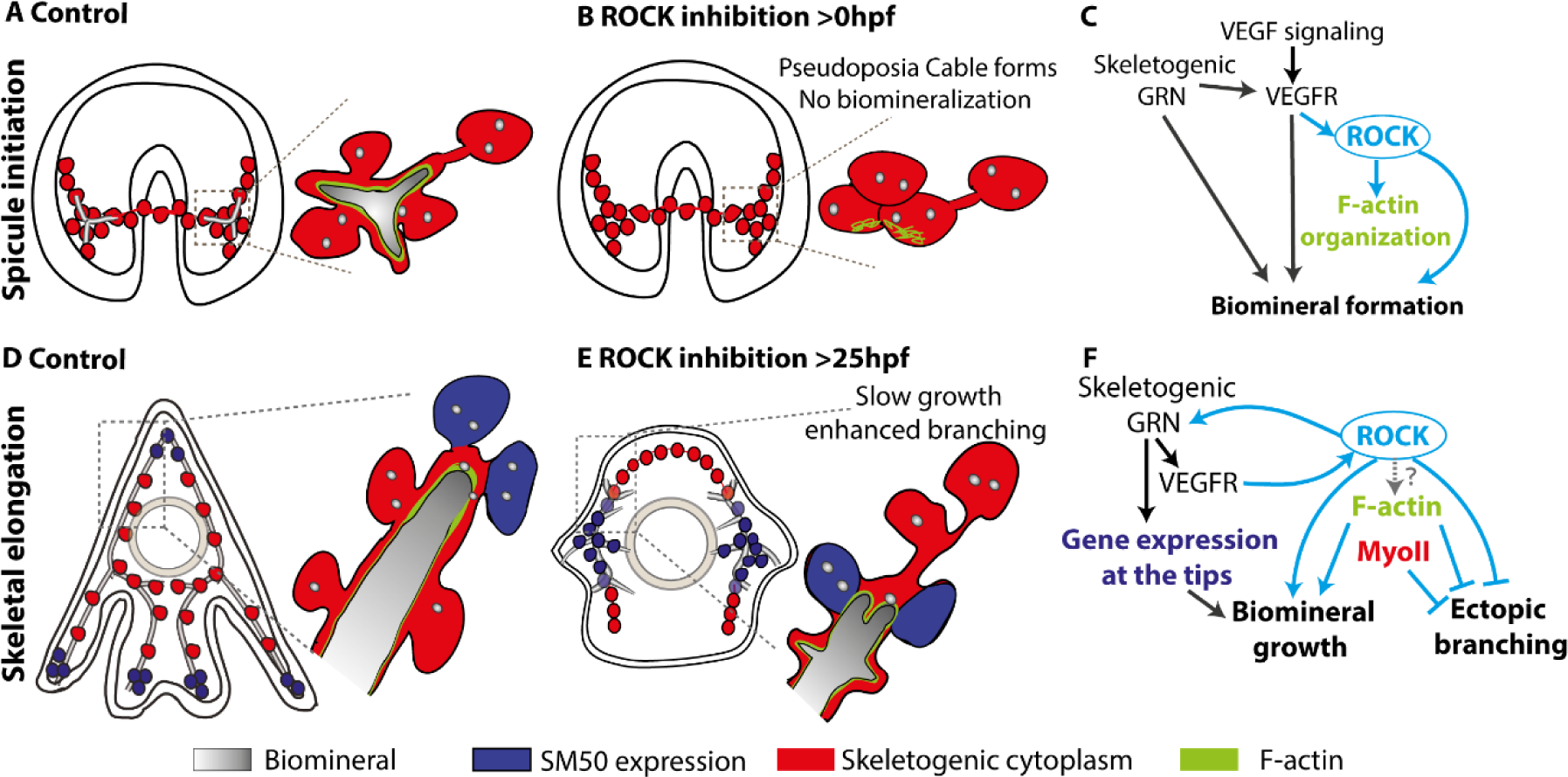
The role of ROCK and the actomyosin network in sea urchin skeletogenesis, summary. (A) the spicule forms at about 24hpf in *P. lividus* and at 27hpf, the triradiate spicule is coated with F-actin. (B) ROCK activity is required for spicule initiation and for F-actin organization around the spicule. (C) model of the functional links between the skeletogenic GRN, ROCK, F-actin and their skeletogenic outcomes, during spicule initiation (∼24-27hpf in *P. lividus*). Cian arrows indicate links discovered in this work. (D) during skeletal elongation F-actin is detected around the spicule cavity and is enriched at the tips of the rods. The expression of SM50 and some other skeletogenic genes, is localized to the tips of the rods. (E) under the addition of ROCK inhibitor, skeletal growth rate is reduced and ectopic spicule branching is observed. The expression of skeletogenic genes is localized to the vicinity of the growing rods. (F) the functional links between the skeletogenic GRN, ROCK, F-actin and MyoII and their skeletogenic outcomes during skeletal elongation. Cian arrows indicate links discovered in this work.

Both continuous and late ROCK inhibition strongly affect the spatial expression of key regulatory and biomineralization genes that are normally expressed differentially within the skeletogenic lineage at the pluteus stage (Figs. 7D-F, S8). These include, the VEGF receptor (VEGFR), the biomineralization genes *SM50* and *SM30,* skeletogenic transcription factors, Ets1, Alx1 and Erg1 that are essential for skeletogenic cell specification and biomineralization (23, 66) and the transcription factors Hex and MyoD1. In normal plutei these genes are differentially expressed within skeletogenic lineage, but in both continuous and late ROCK inhibition the expression of these genes becomes localized to the two lateral skeletogenic cell clusters and their vicinity (Fig. 7D-F, S8). The lateral skeletogenic cells clusters are the cells where these genes are expressed in the gastrula stage (24-27hpf) and where spicule formation begins in normal embryos (6, 32). The expression of VEGFR and SM50 under ROCK inhibition is more expanded than the other tested genes and is also observed at the anterolateral rods (Fig. 7D, E). The VEGF ligand that is normally expressed in the ectodermal cells near the four tips of the post-oral and anterolateral rods (32), is expressed at four ectodermal domains near the lateral skeletogenic cell clusters and near the end of the anterolateral chain, under ROCK inhibition (Fig. S8G). Our data shows that under ROCK inhibition, the expression of key skeletogenic genes remains localized to the area near the lateral skeletogenic cells clusters (Fig. 7D-F, S8), despite the proper formation of the dorsal, anterolateral and mid-ventral chains (Fig. 7C). Overall, ROCK activity is essential for normal expression level and spatial gene expression in the skeletogenic lineage.

## Discussion

Biomineralization is a complex morphogenetic process, encoded in the species genome and executed by the GRNs and the cellular machinery. Here we used the sea urchin larval skeletogenesis to investigate the role of ROCK and the actomyosin network in biomineral formation and in regulating gene expression, as we summarize in Fig. 8. ROCK expression is enriched in the skeletogenic cells, partially depending on VEGF signaling (Fig. 1, 8C). ROCK activity in the skeletogenic cells is necessary for spicule formation, skeletal elongation and prevention of ectopic skeletal branching (Figs. 2-3, S4, S5). Direct inhibition of F-actin polymerization results in similar but not identical skeletogenic phenotypes (Fig. 4). F-actin is organized around the spicule cavity and enriched at the spicule tips (Figs. 5, 6). ROCK activity feeds back into the skeletogenic GRN and affects regulatory and biomineralization gene expression (Fig. 7). Below we discuss our findings and their implications on the biological control and evolution of biomineralization.

The first step in generating the sea urchin spicules is the construction of the spicule cavity where the mineral is engulfed in a membrane coated with F-actin (Fig. 8A). ROCK activity is necessary for the spicule initiation and for the organization of F-actin around the spicule cavity (Figs. 2, 5, 8B). However, since the spicule doesn’t form under ROCK inhibition, it is hard to conclude if the absence of F-actin organization around the spicule is due to ROCK regulation of actin polymerization or due to the absence of spicule in this condition. Relatedly, a direct perturbation of F-actin polymerization strongly disrupts spicule growth and shape, but not spicule formation, indicating that F-actin is not essential for spicule initiation (Fig. 4). We therefore propose that ROCK activity is necessary for spicule initiation, but not through its’ direct regulation of the actomyosin network, but though its other targets, yet to be identified (Fig. 8C).

After the spicule cavity has formed, the spicules elongate inside the pseudopodia chord by mineral deposition at the tips of the spicule rods, which depends on ROCK activity, on F-actin polymerization and to a lesser extent, on MyoII contractility (Figs. 2-4, 8D-F). F-actin is detected along the spicules and is significantly enriched at the tips of the elongating spicules, independently of ROCK activity (Fig. 5, 6, 8D, E (67)). Under ROCK inhibition and under the inhibition of F-actin polymerization, ectopic branching is observed at the tips and in the back (Figs. 2H, J, O, 3A, I, 4, 8E). MyoII inhibition results with ectopic branching at the tips of the rods (Fig. 4B, G), that is similar to the effect of late ROCK inhibition at low concertation (Fig. 2H, K). These correlative similarities between ROCK and the actomyosin perturbations lead us to the following speculations: the low dosage of late ROCK inhibition is perturbing mostly ROCK activation of MyoII contractility while the higher dosage affects factors that control actin polymerization (Fig. 8F). Further studies in higher temporal and spatial resolution of MyoIIP activity and F-actin structures in control and under ROCK inhibition will enable us to test this.

The participation of ROCK, F-actin and MyoII in polarized growth and vesicle exocytosis has been observed in both animals and plants (67, 68). F-actin is used for short range myosin trafficking of vesicles toward the secretion site (67, 69). Once the secretory vesicle fuses with the membrane, it is coated with F-actin that stabilizes the vesicle and enables MyoII binding and ROCK-regulated contraction, necessary for content secretion (48, 49). In plant cells, F-actin accumulate at the growing tip of the cell and assists vesicle exocytosis necessary for polarized tip-growth (68). ROCK, F-actin and MyoII could be regulating the localized exocytosis of mineral-bearing vesicles during sea urchin skeletogenesis. The reduction of spicule growth rate and the enhanced branching could be due to impaired and misplaced vesicle trafficking and deposition under ROCK and F-actin inhibition. Detailed investigation of the kinetics and deposition of mineral-bearing vesicles and the role of actomyosin network in these processes will illuminate the regulation of vesicle exocytosis during biomineralization.

ROCK activity is necessary for correct spatiotemporal expression of regulatory and biomineralization genes (Fig. 7). ROCK inhibition downregulates the expression level of skeletogenic genes (Fig. 7A) and prevents the spatial progression of key skeletogenic genes toward the edges of the skeletal rods (Fig. 7, S8, 8E). This implies that ROCK activity provides regulatory cues to the skeletogenic GRN required for the dynamic progression of gene expression within the skeletogenic lineage. In other systems, ROCK activity was shown to regulate gene expression through various intracellular pathways (70–72). Furthermore, ROCK has an important role in mechano-sensing of matrix rigidity and transducing it into changes in gene expression (73). Relatedly, stiff substrate activates ROCK in pre-osteoblastic cells *in-vitro*, which activates Erk that leads to osteogenic differentiation and upregulation of osteogenic gene expression (72). Sea urchin ROCK could be a part of the mechano-sensing mechanism that detects the high spicule stiffness and transduces it into gene expression, providing a necessary feedback between spicule elongation and the skeletogenic GRN.

Overall, our findings together with the role of the actomyosin network in biomineralization, from calcifying and silicifying single cells to vertebrates’ bones and teeth (15–22, 72), suggest that this molecular machinery is a part of the common molecular tool-kit of biomineralization. Most likely, the actomyosin network was employed independently, downstream of distinct GRNs across Eukaryotes, to control mineral growth and morphogenesis.

## Methods

### Animal and embryos

Adult *Paracentrotous lividus* were obtained from the Institute of Oceanographic and Limnological Research (IOLR) in Eilat, Israel. Eggs and sperm were obtained by injection of 0.5M KCl solution into the adult sea urchin species. Embryos were cultured at 18°C in 0.2µ filtered ASW.

### Imaging

Embryonic phenotypes and WMISH were imaged by Zeiss Axioimager 2. Fluorescent markers were imaged using Nikon A1-R Confocal microscope. All images were aligned in Adobe photoshop and Illustrator.

### VEGFR inhibitor, Axitinib (AG013736) treatment

Axitinib (AG013736, Selleckchem, Houston, TX, USA) was applied as described in (32)

### Western Blot

Embryo cultures treated with 150 nM Axitinib, or DMSO as control, were collected at 22hpf, 27hpf and 33hpf by centrifugation of 10,000g for 2 minutes. We lysed the pellet in 150µL lysis buffer (20 mM Tris-HCl, 150 mM NaCl, 1% Triton X-100, pH 8) with protease inhibitors (protease inhibitor cocktail w/o metal chelator; Sigma, P8340; 1:100) and phosphatase inhibitors (50 mM NaF, 1 mM Na3VO4, 1 mM Na4P2O7, 1mM, βGP) as described in (74). 40µg protein were loaded on 8% SDS-acrylamide gel, transferred to PVDF membranes for Western blot analysis. Anti-ROCK antibody (ab45171, Abcam) was used in a 1:300 dilution followed by anti-rabbit HRP secondary antibody diluted to 1:2500 (Jackson ImmunoResearch 111-035-003). For the loading control, the membrane was incubated with anti-β-tubulin antibody (1:5000; Sigma) followed by anti-mouse antibody 1:5000 (Jackson ImmunoResearch 115-035-003). Quantification of ROCK signal was done using Image studio lite vr. 5.2. ROCK signal was divided by Tubulin signal and then the ratio between ROCK/Tubulin in control vs. VEGFR inhibition was calculated. The graph in Fig. S2 was generated based on three biological replicates (different sets of parents) at 27hpf and 33hpdf and four biological replicates at 22hpf.

### Immunostaining procedure

Phalloidin labeling, MyoIIP (p20-MyosinII), ROCK (anti ROCK2+ROCK1 antibody [EP786Y], AB-ab45171, Abcam), and 6a9 immunostaining were done similarly to (35).

### Quantification of Anti-ROCK and Phalloidin signal

A graphical program was written to allow manual quantification of the fluorescent signal from the spicule regions by identifying “stained” pixels per selected area. The ratio of stained anti-ROCK to region total area was compared between ectodermal and skeletogenic cells, and between control and VEGFR inhibited embryos at 22, 27 and 33hpf. The average ratios and ztest for significance difference from 1, are 22hpf control, ∼1.3, z=0.0006; VEGFR inhibition ∼1, z=0.5, 27hpf control ∼1.8, z<10-11; VEGFR inhibition ∼1.6, z<10-8, 33hpf control ∼4.6, z<10-23; VEGFR inhibition ∼1.5, z<10-5. For phalloidin analyses, embryos at 33hpf of control and ROCK inhibitor addition at 25hpf were used and for each image, two regions were selected: an area at the tip of the spicule and an area at the back of the spicule, about ∼10 microns apart. The ratio of stained phalloidin to region total area was compared between tip and back, in control and ROCK inhibited embryos. Paired t-test was used to compare the differences in phalloidin signal between the tip and back of the spicules across all groups and between control and ROCK inhibited embryos.

### ROCK MASOs injections

Microinjections were conducted similarly to (32). The eggs were injected with an injection solution containing 0.12M KCl, 0.5µg/µl Rhodamine Dextran and 900µM of MASOs. Translation MASOs were designed and synthesized by Gene Tools, Philomath, OR, according to ROCK sequence (40). The sequence of ROCK MASO-1 (5’-AGACATATTTGGAGCCGA[CAT]CCTG-3’) matches the start of translation region and ROCK MASO-2 (5’-TCTCTTGCGTTATATTCCACTAAGT-3’) matches the 5’ region upstream of the start of translation (40). Control embryos were injected in the same concentration with Random MASO. Injected embryos were cultured at 18°C, imaged and scored at 2dpf.

### ROCK inhibitor (Y27632) treatment

Y27632 stock (10005583 CAS Regisrty No. 129830-38-2, Cayman chemical), was prepared by reconstituting the chemical in DMSO. The embryos were treated with Y27632 at a final concentration between 30-80µM, as mentioned in the results. Throughout the paper, control embryos were cultured in DMSO at the same volume as Y27632 solution and no more than 0.1% (v/v).

### ROCK inhibition in isolated skeletogenic cell culture

Skeletogenic cell culture was performed as described in (37) with minor changes. Isolated skeletogenic cells were cultured in CultureSlides (4 chambers polystyrene vessel tissue culture treated glass slide REF 354104) with ASW (Red Sea Fish LTD) containing Gentamicin and Penicillin-Streptomycin (GPS) at 18°C. Media containing 4% (v/v) horse serum (Sigma-Aldrich H1270) in ASW+GPS was added to the culture. ROCK inhibitor, Y27632, was added to the cell culture at 16hpf (before spicule initiation) or 48hpf (after spicule initiation), and images were taken at 72hpf. Each treatment was conducted in three biological repeats except from the >48hpf addition of 120µM which was done in two biological repeats.

### Quantification of skeletal length and statistical analysis

Skeletal length measurement was done as described in (32). The measurements were repeated for three biological repeats for control, ROCK inhibition with 30μM and 80μM, Latrunculin-A 2nM, Blebbistatin 2µM, and 2 biological repeats for 120μM. In ROCK inhibition experiments, a total of 275 skeletons were measured for control, 105 skeletons for 30μM, 122 skeletons for 80μM and 80 for 120μM. In Latrunculin-A and Blebbistatin experiments, a total of 116 skeletons were measured for control, 149 for Latrunculin-A and 107 for Blebbistatin. The data were analyzed in Excel and the statistical analysis was performed using Kruskal-Wallis non-parametric test in Excel as described in (75) and SPSS statistics 27.

### Spicule preparation for micro-computed tomography (SR-µCT) measurement

Control embryos and embryos where 40µM Y27632 was added at 25hpf, were collected by centrifugation of 50mL Falcon tubes at 2000rpm, at 48h and 72hpf. Every tube was washed in 10 mL cold distilled water (DW), transferred into epi-tube and washed three times in 1mL cold DW (incubation at 4°C between every wash, centrifuge at 2000rpm in room temperature for 2min). Then the skeletons were washed three times in 3% NaOCl followed by three washes in DW, one wash in 70% EtOH and one wash in Acetone. Skeletal spicules were dried at room temperature and stored at -20 °C.

### Synchrotron and lab-based SR-µCT

Dry and loose skeletal spicules prepared as described above, were used for the SR-µCT analysis. We analyzed 14 sets of spicules from three (control) and four (ROCK inhibited) pairs of parents, divided into four groups: 2dpf, control, n=44; 2dpf ROCK inhibited, n=93; 3dpf control, n=51; 3dpf ROCK inhibited, n=93, see Table S1). Spicules were glued to sharpened toothpick tips, either in bulk or individually when picked by hand by means of a brush bristle. Each toothpick loaded with spicules, was mounted on a 3ml vial twist-off lid and for transportation securely stored in the vial (Fig. S5A, B). For each set 2-3 µCT-samples were prepared, which allowed the acquisition of statistically relevant sample sizes (40+ spicules/group, Table S1). Tomographic data sets were acquired using the following scanning parameters; for the lab based µCT (RX Solutions, Chavanod, France) 90kV, 143-160µA, 1440 images, average frames 10, frame rates 0.4-1, voxel sizes: 0.45-59µm; and for the synchrotron radiation µCT (Anatomix, Synchrotron SOLEIL, Gif-sur-Yvette, France) 2048 images, angle between projections 0.09°, voxel size: 0.65µm. Tomographic data reconstruction was performed with commercial (RX Solutions, Chavanod, France) and PyHST2 software (A. Mirone et al., Nucl. Instrum. Meth. B 324 (2014) 41-48, doi:10.1016/j.nimb.2013.09.030) at the MPICI and at Anatomix, SOLEIL, respectively.

### Three-dimensional (3D) data analysis

Data visualization, pre-processing including cropping and image type conversion to 16bit (of the Sr-µCT data), and user-augmented segmentation of intact spicules were performed in Amira using off-the-shelf modules (Fig. S5C-E, (76)). Volume and area measurements were performed on the segmented spicules, *i.e.* the label-fields using Amira’s material.statistics module (Volume3D and Area3D). Average spicule thickness and total length measurements were performed with python code, utilizing available python libraries (*e.g.*, skimage, skan, scipy). The code was applied to 3D-tif files created from Amira’s label-fields, which contained the segmented spicules. For length measurements, the spicules were skeletonized and the chord lengths of the branches forming a spicule-skeleton were summed up to obtain the total spicule length. For thickness measurements, we performed Euclidean distance transforms for each spicule using scipy’s ndimage.morphology.distance_transform_edt package, and calculated the average spicule thickness based on the distance measurements obtained at each voxel from spicule-skeleton to the nearest spicule surface. Quality control of our measurements was performed with open source software, Fiji and available plugins (e.g. BoneJ, FiberJ, DiameterJ).

### Calcein staining

A 2mg/ml stock solution of blue calcein (M1255, Sigma) and a 2mg/ml stock solution of green calcein (C0875, Sigma) were prepared by dissolving the chemicals in distilled water. Working solution of 250μg/ml and 150μg/ml, respectively, was prepared by diluting the stock solution in artificial sea water. Blue calcein was added to the embryo culture at 29hpf for one hour and then washed. At 44hpf the green calcein was added for one hour and washed. For FM4-64 staining A 100µg/ml stock solution of FM4-64 (T13320, Life technologies, OR, USA) was prepared by dissolving the chemical in distilled water. Working solution of 2.5 µg/ml was prepared by diluting the stock solution in artificial sea water. Embryos were immersed in working solution about 10 minutes before visualization.

### Latrunculin-A and Blebbistatin treatments

1mM of Lat-A stock (L12370, Thermo Fisher Scientific) was prepared by reconstituting the chemical in DMSO. The embryos were treated with Lat-A at final concentration of 2 nM. 1mM of Blebbistatin stock (13013-5, Enco) was prepared by reconstituting the chemical in DMSO. The embryos were treated with Blebbistatin at final concentration of 2 µM. For the double inhibition experiments, embryos were treated with Blebb at final concentration of 1.5 µM, and with Lat-A at final concentration of 2 nM.

### WMISH procedure

WMISH was performed as described in (32). List of the primers used to make the WMISH probes is available in (32).

### cDNA preparation for QPCR experiments

For ROCK inhibition experiments total RNA was extracted from >1000 sea urchin embryos in each condition using RNeasy Mini Kit (50) from QIAGEN (#74104) according to the kits’ protocol. DNase treatment on column was done using RNase-Free DNase Set-Qiagen (50) (#79254). RNA was reverse-transcribed using the High Capacity cDNA RT kit, AB-4368814 (Applied Biosystems) following the manufacturer’s instructions.

### Quantitative polymerase chain reaction (QPCR)

QPCR was carried out as described in (32). Complete list of primer sequences used for the experiments in provided there.

## Acknowledgments

We are grateful to SOLEIL for provision of synchrotron radiation facilities and we would like to thank Dr. Mario Scheel, Dr. Timm Weitkamp and Dr. Jonathan Perrin for assistance in using beamline ANATOMIX for nano and µCT. We thank the Bioimaging Unit, Faculty of Natural Sciences, University of Haifa for assistance with the use of confocal. We thank Charles Ettensohn for the generous gift of the 6a9 antibody produced in his lab. We thank David Ben-Ezra, Michael Kantorovitz and Alvaro Israel for their help with sea urchin handling. We thank Haguy Wolfenson for illuminating discussions of the results and conceptual framework. We thank Nir Sapir and Tovah Nehrer for their advice and help with the statistical analysis of spicule length. We thank Aleksei Tabachnic for his help with microinjection. We thank Alvaro Israel and Omri Nahor for their help with sea urchin maintenance. We thank Yulia Kagan for her help with the western blot.

