## Supplementary material for "ROCK and the actomyosin network control biomineral growth and morphology during sea urchin skeletogenesis": Sup figs and tables

**Supplementary material for - Actomyosin remodeling regulates biomineral formation, growth and morphology during eukaryote skeletogenesis**

Eman Hijaze[1], Tsvia Gildor[1], Ronald Seidel[2], Majed Layous[1], Mark Winter[3], Luca Bertinetti[2], Yael Politi[2] and Smadar Ben-Tabou de-Leon[1,*]

[1]Department of Marine Biology, Leon H. Charney School of Marine Sciences, University of Haifa, Haifa 31905, Israel.

[2]B CUBE Center for Molecular Bioengineering, Technische Universität Dresden, 01309 Dresden, Germany.

[3]Department of Electrical Engineering, Computer Science and Mathematics, Technische Universiteit Delft, Delft 2628CD, Netherlands

**This PDF file includes:**

**Figs. S1 to S8**

**Tables S1 to S4**

### Figures

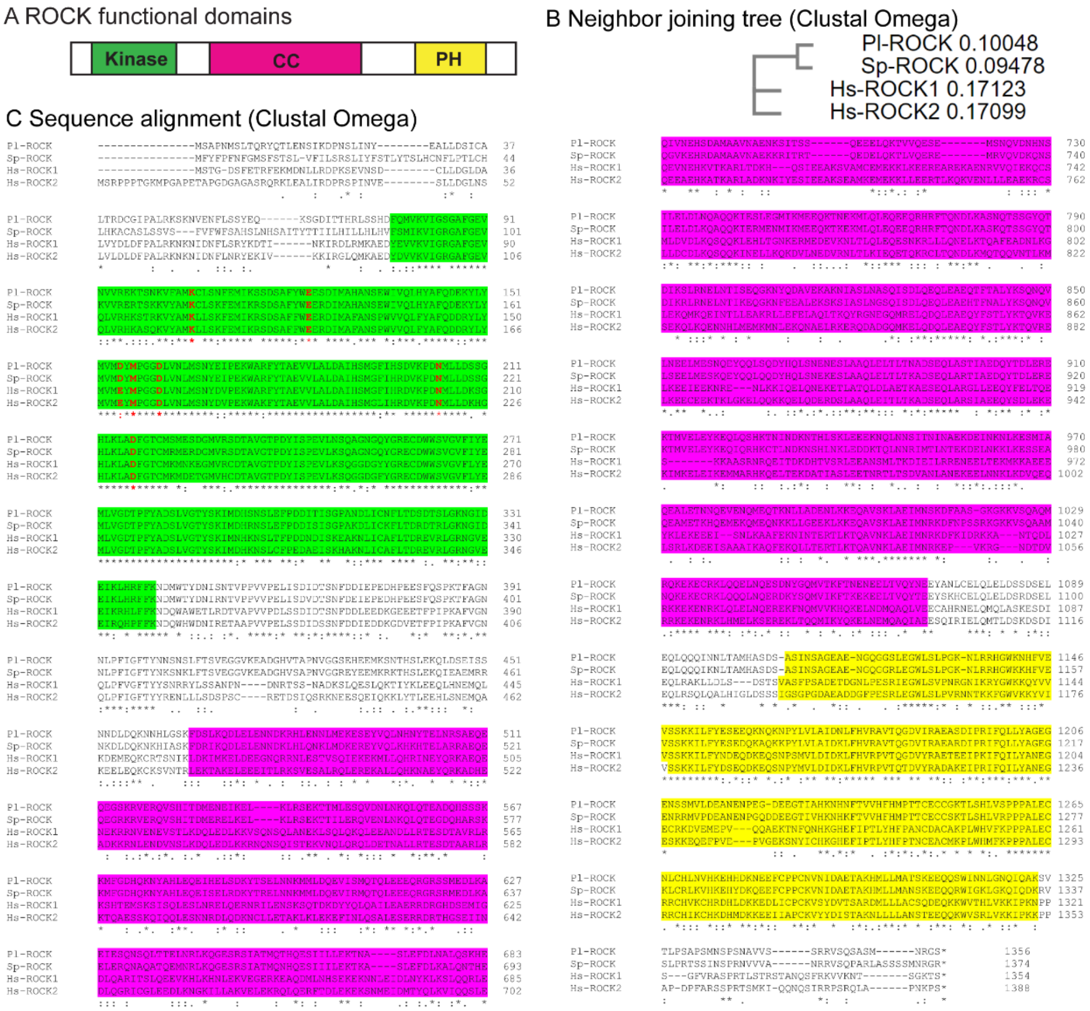


**Sup Figure 1 Sea urchin ROCK sequence and conserved domains.** (A) Diagram showing the functional domains in the human ROCK protein. (B) Neighbor joining tree of human ROCK1, ROCK2 and the sea urchin genes using clustal omega showing unrooted tree. (C) sequence alignment of Pl-ROCK, Sp-ROCK, Hs-ROCK1 and Hs-ROCK2. The amino acids where Y-27632 binds are marked in red (1). Asterisk indicates amino acid identity; highly similar amino acids are indicated in two dots and similar amino acids are indicated in one dot. Six of the Y-27632 bound amino acids are conserved and the 7$^{th}$ one is highly similar between human and sea urchin (E->D). Yellow background indicates the PH domain, green background the kinase domain, Pink background indicates the coil-coil domain.

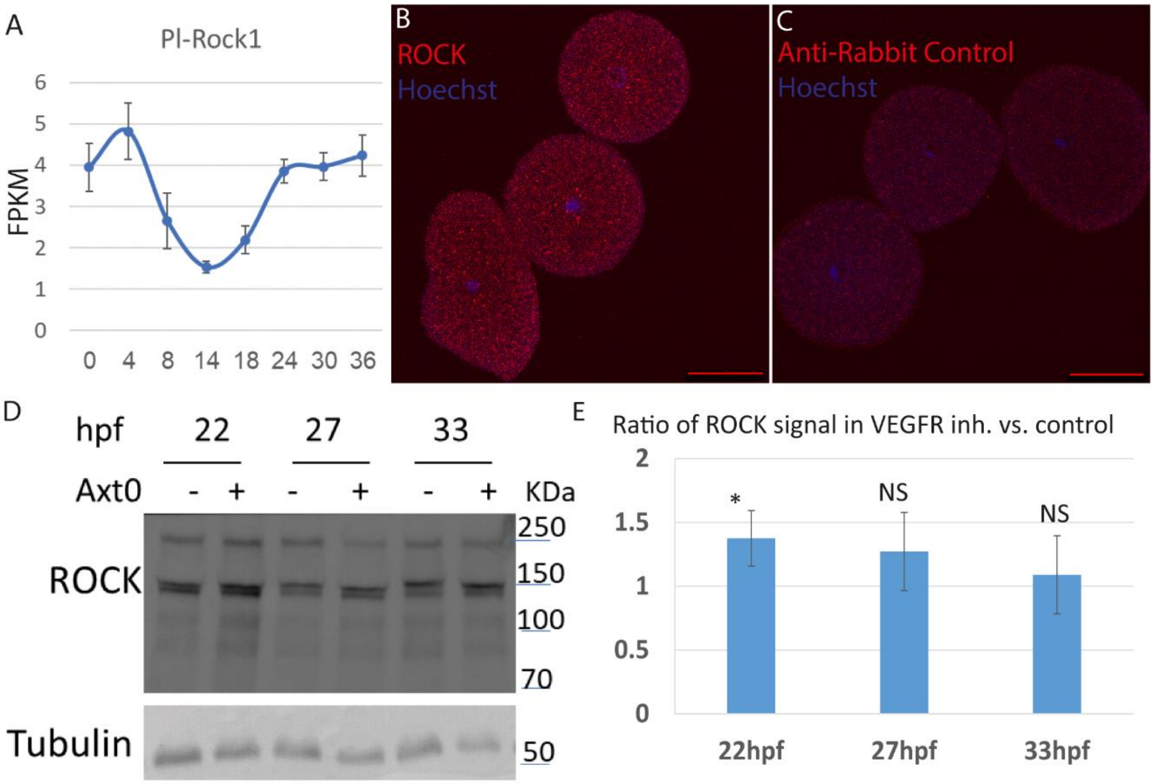


**Sup Figure 2 Time course of *Pl-ROCK*, immunostaining and western blot of ROCK antibody in sea urchin embryos.** (A) Time course of Pl-ROCK measured by RNA-seq in fragments per kilobase of transcript per million mapped reads (FPKM). Error bars indicate standard deviation over three independent biological replicates. The data is taken from Morgulis et al, PNAS 2019 (2). (B) ROCK immunostaining (red) in fertilized eggs, overlaid with Hoescht nuclear staining (blue). (C) Negative control, using only the secondary anti-rabbit antibody (red), overlaid with Hoescht nuclear staining (blue). The experiment was done in three biological replicates, where in ROCK, n=80, and in the negative control n=81 embryos were scored. Scale bars in B and C are 50µm. (D) representative image of western blot analysis of the ROCK antibody using crude extracts of whole embryos at 22hpf, 27hpf and 33hpf, in control and VEGFR inhibited embryos. (E) Quantification of ROCK protein abundance in VEGFR inhibition compared to control embryos at the three time points. ROCK protein abundance was normalized to the Tubulin signal. Statistical significance was measured using z-test where, * indicates $P<0.05$, and NS - non-significant, $p>0.05$ ($p=0.04$ at 22hpf, $p=0.22$ at 27hpf and $p=0.4$ at 33hpf). The experiments were conducted in four biological replicates for 22hpf and three biological replicates for 27hpf and 33hpf.

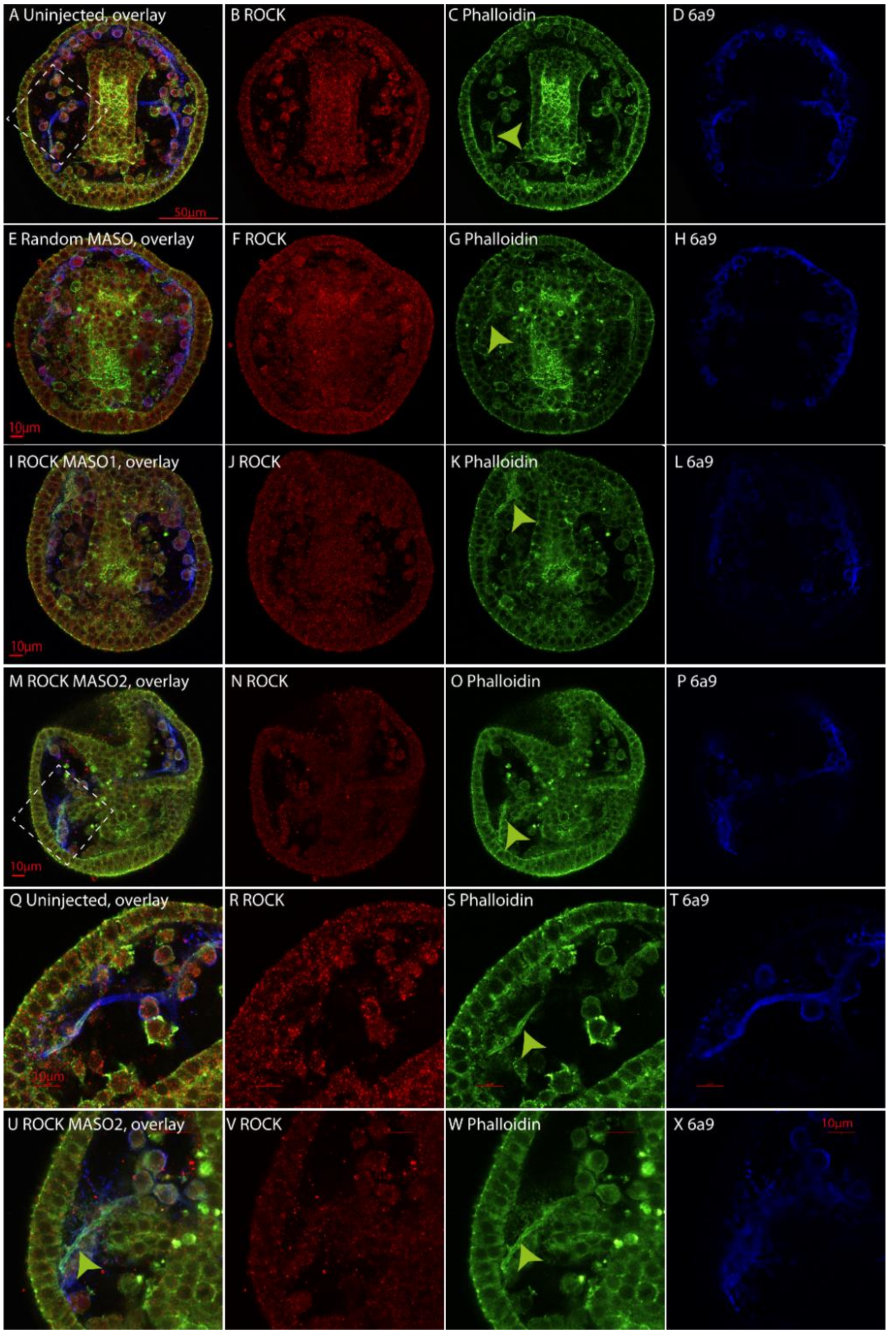


**Sup Figure 3 ROCK and F-actin expression in control and ROCK MASO injections at 33hpf.** Representative images of embryos stained with ROCK antibody (red), Phalloidin (green) and the skeletogenic marker 6a9 (blue) at 33hpf. (A-D) Uninjected control, (E-H) Random MASO, (I-L) ROCK MASO1, (M-P) ROCK MASO2, (Q-T) Enlargement of marked square in A of Uninjected embryo, showing minor overlap between the ROCK and the Phalloidin signal. (U-X) Enlargement of the marked square in M of embryo injected with ROCK MASO2. The green arrows point to the enriched F-actin signal around the spicule. These experiments were done in three independent biological replicates where the total number of embryo scored was n=24 uninjected embryos; n=25 Random MASO; n=35 ROCK MASO1 and n=35 ROCK MASO2. Scale bars are 50µm in A and 10µm elsewhere.

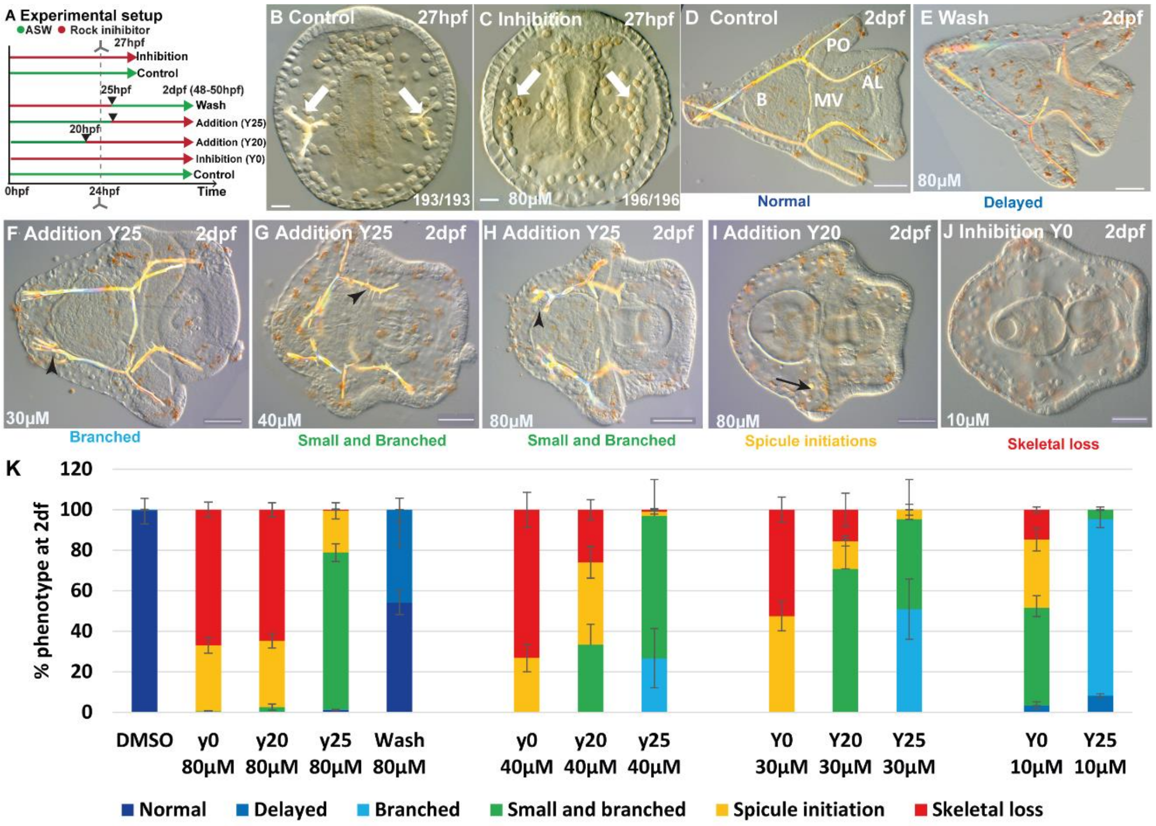


**Sup Figure 4 ROCK inhibition phenotypes for different treatments and concentrations.** (A) experimental design of the various ROCK treatments showing the time of inhibitor addition/wash. (B) representative control embryo and (C) embryo under ROCK inhibition using 80µM of the inhibitor Y27632 at 27hpf. The numbers at the bottom indicate the number of embryos that show this phenotypes (left) over all observed spicules (right). (D-J) various skeletogenic phenotypes due to ROCK inhibition at different times and concentrations observed at 2dpf, organized from normal to most sever skeletogenic phenotypes. Treatment and concentrations are indicated for each image. (K) Summary of ROCK perturbation phenotypes at 2dpf. Color codes of phenotypes match the color of the phenotypes reported in B-H. Scale bars are 20µm in B, C and 50µm in D-J. Results are based on 3-8 biological replicates for the different treatments, exact number of replicates and embryo scores for each experiment is provided in Table S1.

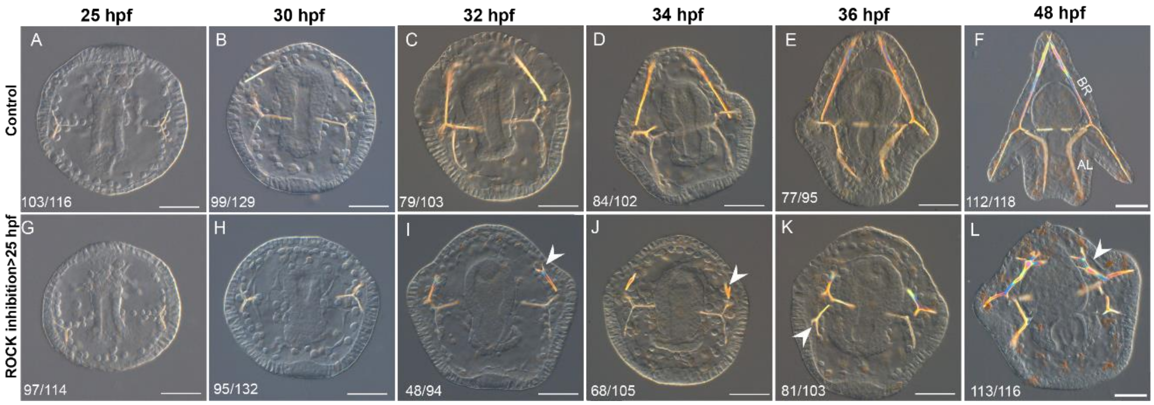


**Sup Figure 5 Time course of the effect of ROCK inhibition after 25hpf.** (A-F) representative images of control embryos from gastrulation (25hpf) until pluteus stage (48hpf). (G-L) representative images of embryos treated with 30µM ROCK inhibitor Y27632 from 25hpf and on, at equivalent developmental stages to A-F. Arrowheads points to ectopic branching. Three biological replicates were conducted for each treatment and the numbers at the bottom left of each panel indicate the number of embryos that show this phenotype out of all embryos scored. Scale bars are 50µm. BR- body rods, AL- anterolateral rods.

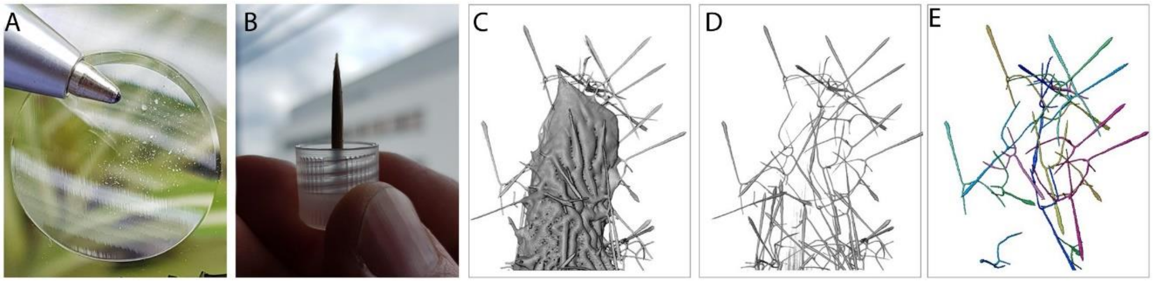


**Sup Figure 6 Sample preparation for SR-µCT data acquisition and analysis.** (A) Dissected, dried spicules were used for tomographic image acquisition. (B) Typically, hundreds to a dozen spicules were glued to sharpened toothpick tips for sample fixation. Tooth picks were mounted on 3ml vial twist-off lids and securely stored in the vial. (C) 3-D rendering of an exemplary tomographic data set, showing multiple calcitic spicules attached to the toothpick tip. (D) User-augmented data segmentation (excluding background and toothpick) and visualization of intact spicules were performed in Amira. (E) Volume, area, length and thickness measurements were performed on segmented and labeled spicules as described in the methods section.

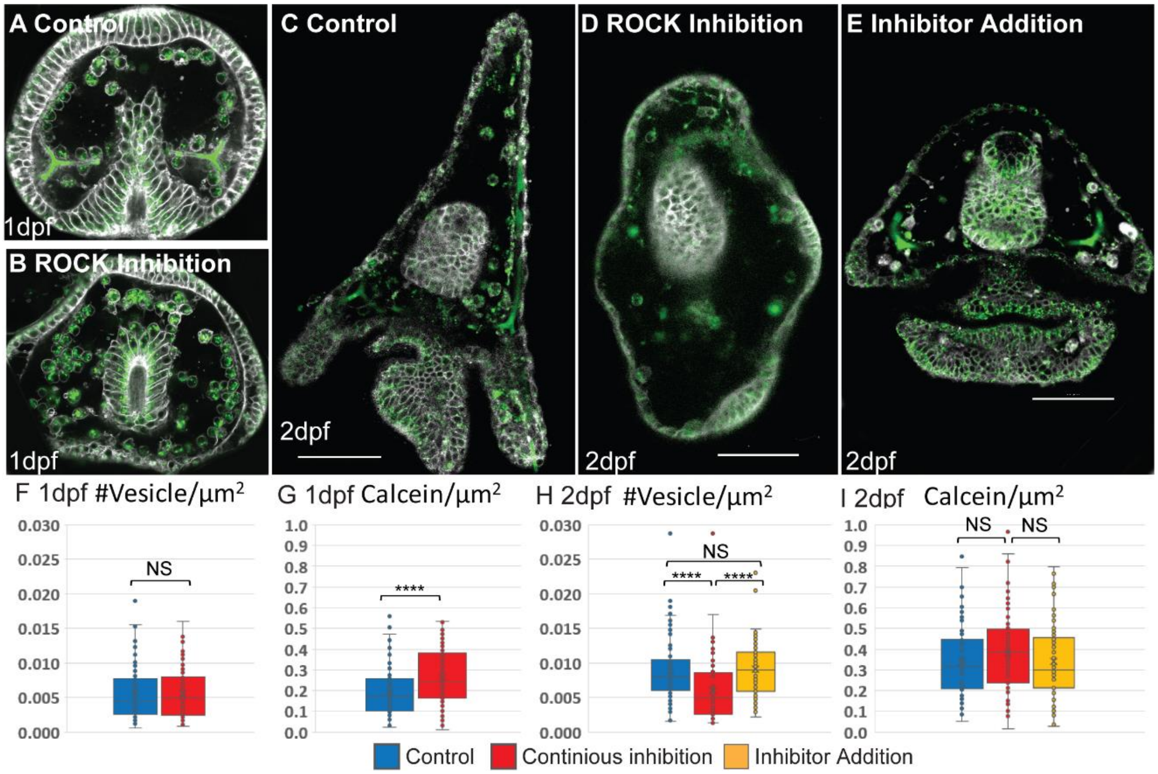


**Sup Figure 7 Effect of ROCK inhibition on calcein stained area and vesicle number.** A-E, confocal images of calcein staining (green) and FM4-64 membrane marker (white) show the presence of calcium vesicles in the skeletogenic cells in normal and ROCK inhibited embryos at 1dpf and 2dpf. Scale bars are 50µm. F, H, vesicle number per $\mu m^2$ in the skeletogenic cells in control and ROCK inhibition at 1dpf (F) and 2dpf (H). G, I, calcein pixels per $\mu m^2$ in the skeletogenic cells in control and ROCK inhibition at 1dpf (G) and 2dpf (I). Inhibition refers to continuous inhibition from fertilization, and addition refers to the addition of the inhibitor at 25hpf. Experiments were performed in four independent biological replicates where in each condition at least 30 embryos were measured. Each box plot shows the median (black line), average (x) the first and the third quartiles (edges of boxes) and all measured points. Statistical significance was measured using paired 2-tailed t-test where, * indicates $P<0.05$, and *** indicates $P<0.0005$. (n=3, exact number of cells in each condition is provided in SI appendix, table S1).

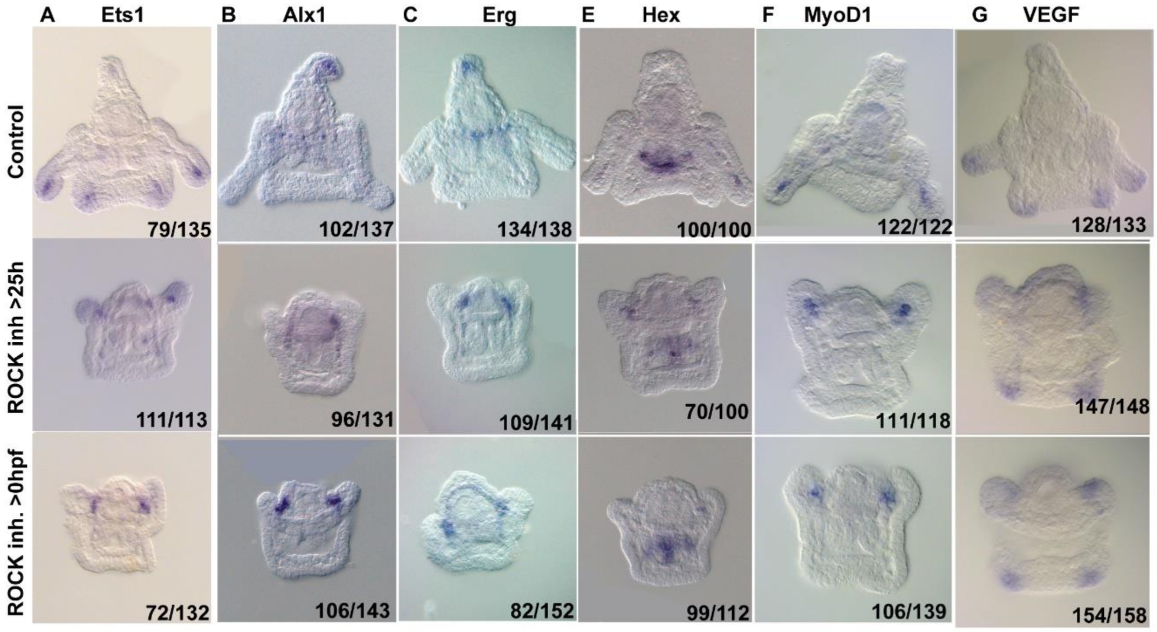


**Sup Figure 8 ROCK activity is essential for normal gene expression in the skeletogenic cells.** Representative images of control embryo (top panels), embryos where ROCK inhibitor was added at 25hpf (middle panels), and embryos that were exposed to continuous ROCK inhibition (bottom panels), at the pluteus stage (~48hpf). Gene names are indicated at the top of each panel. Numbers at the bottom of each image indicate the number of embryos that show this phenotype (left) out of all embryos scored (right), conducted in at least three independent biological replicates.

#### Tables

**Table S1** ROCK inhibition experimental details

| **Condition** | **# replicates** | **# embryos scored** |
|---|---|---|
| DMSO (control) | 8 | 438 |
| Y0 10µM | 3 | 122 |
| Y25 10µM | 3 | 109 |
| Y0 30µM | 3 | 120 |
| Y20 30µM | 3 | 103 |
| Y25 30µM | 3 | 108 |
| y0 40µM | 3 | 93 |
| y20 40µM | 3 | 123 |
| y25 40µM | 3 | 101 |
| y0 80µM | 8 | 377 |
| y20 80µM | 3 | 116 |
| y25 80µM | 7 | 369 |
| Wash 80µM | 5 | 268 |

**Table S2 µCT statistics of control and ROCK inhibited spicules**

| | # of spicules | Avrg. length | Stdev. length | Avrg. Thickness | Stdev. Thickness | Avrg. volume | Stdev. volume | Avrg. area | Stdev. area |
|---|---|---|---|---|---|---|---|---|---|
| **Control_48h** | 44 | 497.2 | 62.43 | 3.7 | 0.44 | 6860.28 | 1999.61 | 5822.46 | 1121.57 |
| **Control_72h** | 51 | 697.4 | 71.69 | 4.4 | 0.37 | 14673.2 | 2747.3 | 9896.75 | 1199.63 |
| **Y25_48h** | 93 | 187.93 | 50.18 | 3.92 | 0.53 | 2925.29 | 973.86 | 2324.51 | 606.62 |
| **Y25_72h** | 94 | 276.29 | 81.8 | 4.46 | 0.65 | 5799.48 | 2408.91 | 4002.78 | 1250.25 |

**Table S3 Statistical significance between μCT measurement of control and ROCK inhibited spicules at 2dpf and 3dpf**

| | **Y25 48hpf** |
|---|---|
| **Control 48hpf** | Length: p=2.2e-63, t=31.0771 |
| | Thickness: p=0.02, t=-2.342 |
| | Volume: p=8.1e-32, t=15.5207 |
| | Area: p=6.3e-50, t=23.6854 |
| | **Y25 72hpf** |
| **Control 72hpf** | Length: p=3.9e-65, t=30.8792 |
| | Thickness: p=0.5, t=-0.6798 |
| | Volume: p=2.2e-43, t=20.1038 |
| | Area: p=1.2e-58, t=27.4415 |

**Table S4 Lantrunculin-A and Blebbistatin experimental details (whole embryos)**

| Treatment | # replicates | # scored embryos |
|---|---|---|
| **DMSO (control)** | 5 | 176 |
| **LatA >20hpf** | 4 | 152 |
| **LatA >25hpf** | 4 | 140 |
| **Bleb >20hpf** | 4 | 162 |
| **Bleb>25hpf** | 4 | 144 |
| **LatA+Bleb>20hpf** | 3 | 96 |
| **LatA+Bleb>25hpf** | 3 | 114 |


1. M. Jacobs *et al.*, The structure of dimeric ROCK I reveals the mechanism for ligand selectivity. *J Biol Chem* **281**, 260-268 (2006).
2. M. Morgulis *et al.*, Possible cooption of a VEGF-driven tubulogenesis program for biomineralization in echinoderms. *Proc Natl Acad Sci U S A* **116**, 12353-12362 (2019).
